# Tissue architecture and immune niches govern ctDNA release in colorectal cancer

**DOI:** 10.1101/2025.11.28.691139

**Authors:** Stefan Kühberger, Katja Sallinger, Christin-Therese Müller, Maria Escriva Conde, Sergio Marco Salas, Silvia Andaloro, Christine Beichler, Ricarda Graf, Karin Pankratz, Julia Enzi, Sarah Binder, Martina Scheiber, Lilli Bonstingl, Jasmin Blatterer, Stefan Uranitsch, Gabriele Moitzi, Hannes Schmölzer, Hubert Hauser, Karin Strohmeyer, Mats Nilsson, Sigurd Lax, Antonia Syrnioti, Rudolf Oehler, Gerald Höfler, Felix Aigner, Amin El-Heliebi, Ellen Heitzer

**Affiliations:** Institute of Human Genetics, Diagnostics and Research Center for Molecular Biomedicine, Medical University of Graz, Graz, Austria; Division of Cell Biology, Histology and Embryology, Gottfried Schatz Research Center for Cell Signalling, Metabolism and Aging, Medical University of Graz, Graz, Austria; Science for Life Laboratory, Department of Biochemistry and Biophysics, Stockholm University, Solna, Sweden; Department of Surgery, St. John of God Hospital Graz, Graz, Austria; Department of Surgery, General Hospital Graz West, Graz, Austria; Department of Surgery, General Hospital Leoben, Leoben, Austria; Department of Pathology, General Hospital Graz West, Graz, Austria; Department of General Surgery, Division of Visceral Surgery, Medical University of Vienna, Vienna, Austria; Institute of Pathology, Diagnostics and Research Center for Molecular Biomedicine, Medical University of Graz, Graz, Austria; European Liquid Biopsy Society (ELBS), Hamburg, Germany

## Abstract

Circulating tumor DNA (ctDNA) is central to liquid biopsy-based cancer detection, yet its release into the bloodstream varies widely and remains poorly understood. To define the tissue-level determinants of ctDNA shedding in colorectal cancer (CRC), we integrated tumor-informed plasma sequencing with detailed histopathology, immunophenotyping, spatial transcriptomics, and *in situ* mutation detection in resectable stages (I–III). ctDNA detectability increased with tumor burden, and high ctDNA shedders exhibited a distinct architectural and microenvironmental phenotype characterized by expanded necrotic pseudolumina, frequent epithelial barrier disruption, and dense myeloid infiltration. Spatial profiling revealed stress-associated malignant programs and a myeloid-rich immune-luminal niche. *In situ* sequencing confirmed plasma-detected mutations within pseudoluminal debris, identifying these structures as focal reservoirs of shed DNA. These findings provide a mechanistic framework linking tissue architecture, immune remodelling, and spatially organized cell death to ctDNA release with implications for refining liquid biopsy applications.

## Introduction

Circulating tumor DNA (ctDNA) is a tumor-derived fraction of cell-free DNA (cfDNA) in the bloodstream and represents a cornerstone of liquid biopsy applications for cancer detection, disease monitoring, and treatment guidance (1–3). While ctDNA levels generally decline after surgery and are higher in advanced-stage cancers, detection rates vary widely, even within the same stage and tumor type (9–11), with 10–40% of late-stage cancers showing no detectable ctDNA with conventional sequencing technologies (4–7). This variability reflects biological determinants of ctDNA generation, release, and clearance that remain poorly understood (4,5,8). While stage, tumor burden, and molecular features explain part of the variation (9–13), the contributions of histopathology, tumor microenvironment, and mechanisms of cell death are less explored. Moreover, patient-specific factors such as age, comorbidities, inflammatory states, and tissue injury may be key determinants of ctDNA levels (14–17). These factors modulate cell death and turnover in tumor and normal cells, especially those of hematopoietic, epithelial, and endothelial origin, thereby affecting ctDNA detectability. The limited understanding of how, when, where and why ctDNA enters the circulation represents a fundamental barrier for the development of blood-based early cancer detection tests, where ctDNA is present only in trace amounts and is heavily diluted by cfDNA from normal cells. Without a mechanistic framework describing the earliest stages of ctDNA shedding during tumorigenesis, it is challenging to identify optimal molecular targets or refine analytical strategies to maximize assay sensitivity and specificity.

Colorectal cancer (CRC), one of the most common cancers worldwide, offers a clinically relevant model for dissecting these mechanisms. Early detection improves survival, yet current screening uptake is suboptimal, and serum protein markers lack the performance for population screening (18). ctDNA-based assays could provide a sensitive, non-invasive alternative, if we can identify the biological contexts in which tumors shed detectable DNA. In fact, several ctDNA-based applications in CRC have already demonstrated clinical utility, including detection of minimal residual disease (MRD) (19), monitoring of treatment response (20), and identification of actionable mutations and resistance mechanisms (21). Importantly, CRC is considered a relatively high-shedding tumor type compared to many other solid malignancies (4,5,22,23), making it particularly well suited to investigate the tissue and microenvironmental features that may promote ctDNA release.

In this study, we combined ctDNA detection rates with detailed histopathological, immunohistochemical (IHC), spatial genetic and transcriptomic profiling of resectable CRC (stages I–III) to define tissue-level determinants of ctDNA shedding. To this end, we applied a matched panel tumor-informed ctDNA detection that balances the biological specificity of bespoke assays with the broader, unbiased coverage of tumor-agnostic designs and is therefore ideal for mechanistic studies of ctDNA release.

By correlating ctDNA presence and levels with detailed tissue characteristics, we systematically evaluated the interplay between tumor biology, microenvironmental factors, and ctDNA shedding. This integrated approach provides one of the most granular datasets to date on the determinants of ctDNA shedding in solid tumors and offers a framework for refining the design of next-generation, blood-based early detection assays in CRC and beyond.

## Results

### ctDNA shedding correlates with stage and tumor size

To quantify treatment-naïve ctDNA levels with high analytical sensitivity, a total of 59 patients (stage I, *n* = 17; stage II, *n* = 22; stage III, *n* = 20) underwent matched targeted sequencing of resected primary tumor tissue and pre-surgery plasma (***Table 1, Fig. 1A***). To ensure a comprehensive and unbiased representation of tumor-derived variants in the circulation, we applied the same broad genomic profiling assay (TruSight Oncology 500, Illumina) to both tissue and plasma samples. This harmonized approach captures all relevant tumor variants while restricting plasma analysis to mutations identified in the matched tumor, thereby reflecting which tumor-derived DNA molecules are more abundant in the circulation.

**Figure 1.**
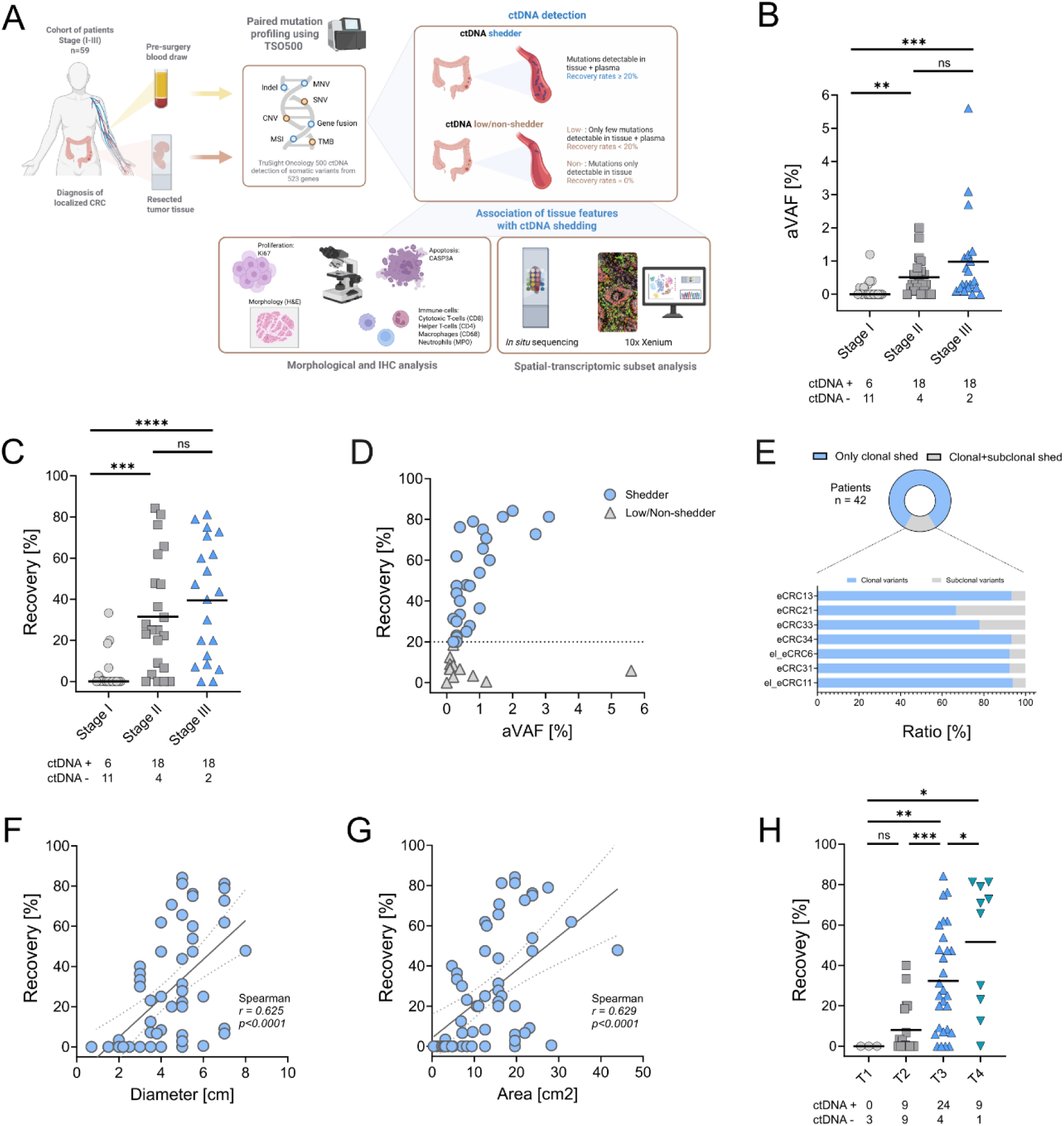
ctDNA shedding correlates with size and stage in localized CRC. **A)** Schematic study design including paired tumor-plasma sequencing (TSO500), ctDNA detection and consequent correlation with IHC and spatial transcriptomics analyses of tumor architecture and microenvironment. **B)** Average variant allele frequencies (aVAF) and **C)** primary tissue recovery rates in plasma across pathological stages I-III. Lines indicate mean values of all patients, *p-*values were calculated using two-tailed Mann–Whitney U test. **D)** Dot-plot showing association of aVAFs with recovery, dotted line indicates cut-off value for ctDNA low/non-shedder at 20% recovery **E)** Top: Proportion of ctDNA positive patients showing either clonal or joint clonal and subclonal variants in plasma, Bottom: Relative contribution of clonal and subclonal variants in 7 patients that show shedding of subclonal variants. **F)** Association of tumor diameter and **G)** tumor area with recovery. *r*- and *p*-values were calculated using Spearman rank-correlation, a linear regression line with 95% confidence interval is shown for visual reference. **H)** Recovery rates across T-stage, *p-*values were calculated using two-tailed Mann–Whitney U test.

**Table 1:** Clinical Correlates of ctDNA shedding.

|  | **SHEDDER**<br>**(N=31)** | **LOW/NON-SHEDDER**<br>**(N=28)** | **P-VALUE** |
| --- | --- | --- | --- |
| AGE (MEAN+ SD) | 68.5 (+ 12.3) | 73.3 (+ 8.6) | 0.086 |
| **SEX** |  |  |  |
| FEMALE | 11 | 11 | 0.793 |
| MALE | 20 | 17 | 0.793 |
| CEA- PREOPERATIVE [NG/ML] | 8.5<br>(+9.2; 1.7-74.5) | 5.6<br>(+ 5.1; 1.5-39.6) | 0.682 |
| MSI | 4 | 9 | 0.116 |
| TMB | 15.4<br>(+15.5; 2.1-130.6) | 32.1<br>(+30.5; 2.3-121.2 ) | 0.108 |
| BRAF V600E POS. | 4 | 6 | 0.494 |
| KRAS MUTATED | 9 | 10 | 0.781 |
| **LOCALIZATION PRIMARY** |  |  |  |
| CECUM | 0 | 2 | 0.221 |
| C.ASCENDENS/CECUM | 4 | 1 | 0.356 |
| C.ASCENDENS | 8 | 10 | 0.572 |
| C.TRANSVERSUM | 1 | 4 | 0.18 |
| C.DESCENDENS | 1 | 3 | 0.337 |
| C.SIGMOIDEUM | 12 | 3 | **0.018\*** |
| C.SIGMOIDEUM/RECTUM | 2 | 1 | 1.000 |
| RECTUM | 3 | 4 | 0.698 |
| **T-STAGE** |  |  |  |
| T1 | 0 | 3 | 0.101 |
| T2 | 4 | 14 | **0.004\*\*** |
| T3 | 19 | 9 | **0.037\*** |
| T4 | 8 | 2 | 0.084 |
| **N-STAGE** |  |  |  |
| N0 | 17 | 22 | 0.097 |
| N1 | 9 | 3 | 0.110 |
| N2 | 5 | 3 | 0.710 |
| **UICC-STAGE** |  |  |  |
| 1 | 2 | 15 | **<0.001\*\*\*** |
| 2 | 15 | 7 | 0.105 |
| 3 | 14 | 6 | 0.097 |
| **GRADE** |  |  |  |
| LOW GRADE (G1-G2) | 27 | 24 | 1.000 |
| HIGH GRADE (G3-G4) | 4 | 4 | 1.000 |
| **L - LYMPHATIC INVASION** |  |  |  |
| L0 | 6 | 9 | 0.479 |
| L1 | 10 | 7 | 0.479 |
| NA | 15 | 12 |  |
| **V - VENOUS INVASION** |  |  |  |
| V0 | 9 | 11 | 0.476 |
| V1 | 1 | 0 | 0.476 |
| NA | 21 | 17 |  |
| **TUMOR BUDDING** |  |  |  |
| YES | 4 | 2 | 0.061 |
| NO | 0 | 5 | 0.061 |
| NA | 27 | 21 |  |

Since an initial validation demonstrated no significant difference in sequencing performance between fresh-frozen and FFPE colorectal cancer tissue (***Supplementary Fig. 1***), all subsequent tissue analyses were conducted using FFPE material. The mutational profiles reflected expected CRC landscapes, with *TP53* and *APC* being among the most frequently mutated genes (***Extended Data Fig. 1***). At least one tumor-specific variant was identified in cfDNA of 6/17 (35.2%) stage I, 18/22 (81.8%) stage II, and 18/20 (90.0%) stage III patients. Mean plasma variant allele frequencies (aVAF) were 0.2% in stage I, and significantly increased to 0.6% in stage II, and 0.8% in stage III, respectively (***Fig. 1B***). Given that low-level VAFs are highly susceptible to technical and biological variability, they provide only a coarse and often unreliable estimate of the ctDNA fraction. In contrast, ctDNA recovery rates, defined as the proportion of somatic tumor variants detected in plasma, exhibited a similar but more statistically robust stage dependency, with mean values of 13.6% in stage I (range 0.5–33.3), 38.6% in stage II (range 3.4–84.2), and 43.9% in stage III (range 5.9–81.3) (***Fig. 1C***). Moreover, correlation analyses demonstrated substantial dispersion of recovery rates among samples with aVAFs between 0–1%, indicating that recovery more accurately reflects biological heterogeneity at low ctDNA levels and offers greater discriminatory sensitivity in low-shedding contexts (***Fig. 1D***). Hence, patients were divided into ctDNA shedders (recovery rates >= 20%) and a joint group of low/non-shedders (recovery rates < 20%). This threshold-based grouping served as the primary classification scheme used across all subsequent analyses in the study.

Interestingly, neither tumor mutational burden (TMB) nor microsatellite instability (MSI) showed a significant association with recovery rates (***Supplementary Fig. 2, Table 1***). As expected, clonal variants were more consistently detected in plasma, whereas subclonal variants, particularly those at lower tumor allele frequencies, were recovered less frequently, reflecting the biological and analytical challenges of detecting heterogeneous tumor populations in circulation (***Fig. 1E***).

When assessing morphological parameters, both tumor size (diameter and cross-sectional area) and T-stage were significantly correlated with ctDNA recovery, indicating that tumor burden and local invasion jointly influence ctDNA shedding (***Fig. 1F-H***). Nevertheless, substantial inter-patient variability in ctDNA detection persisted within each stage, suggesting that size and invasion depth alone do not fully explain the biological determinants of shedding. Comparative analysis of further clinicopathological parameters including tumor grade, lymphovascular invasion as well as localization revealed no additional significant associations (***Table 1***).

### Cell death-associated architecture, epithelial barrier loss, and immune infiltration in ctDNA shedders

Given that ctDNA shedding into the circulation is likely shaped by complex microenvironmental processes and various cell death modalities (24,25), we performed comprehensive morphological and IHC analyses in resected CRC specimens from 49 of the 59 patients (shedders, *n* = 28; low/non-shedders, *n* = 21) to identify histological features associated with ctDNA shedding status. Ten cases with predominant mucinous architecture were excluded from pseudoluminal analyses, as mucinous areas could not be reliably distinguished from pseudoluminal structures. Quantitative histopathological assessment revealed a striking architectural distinction between shedders and low/non-shedders. Tumors from ctDNA shedders exhibited large, confluent glandular formations with expansive pseudoluminal spaces often filled with necrotic detritus, whereas low/non-shedders displayed more densely packed epithelial areas with narrow or poorly defined lumina (***Fig. 2A***). To control for tumor size and cellular density, the pseudoluminal area was normalized to cytokeratin-positive epithelial area, yielding a pseudoluminal-to-epithelial area ratio that was significantly higher in shedders (median 0.49 vs. 0.16, *p*=0.0002; ***Fig. 2B***).

**Figure 2.**
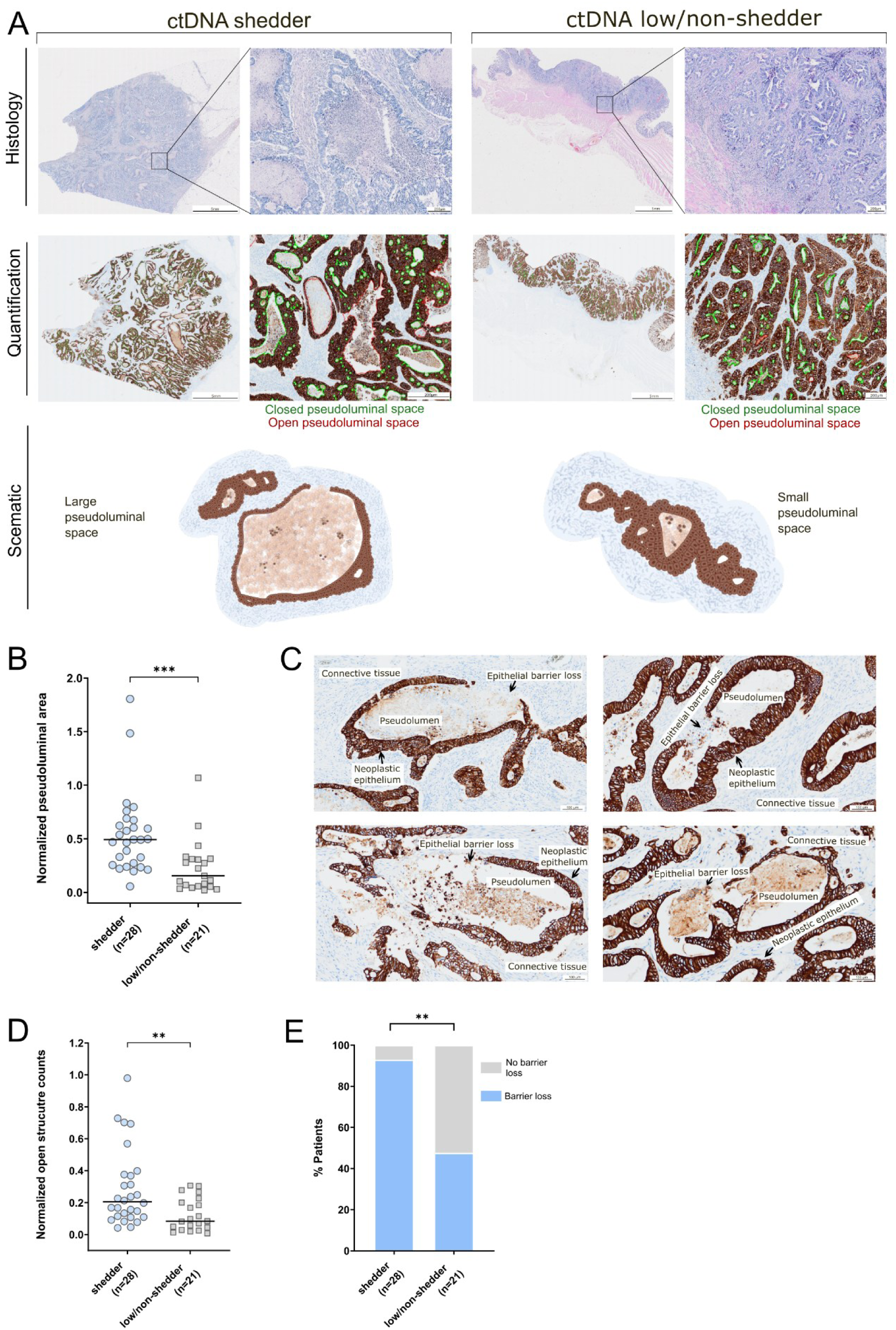
Large pseudoluminal area and epithelial barrier loss is associated with ctDNA shedding in colorectal cancer. **A)** Representative Hematoxylin and Eosin (H&E) and Cytokeratin (CK) immunohistochemistry (IHC) stainings of ctDNA shedders on the left and low/non-shedders on the right. Shedders display large, glandular structures with expanded pseudoluminal areas frequently containing necrotic material, whereas low/non-shedders show compact epithelial arrangements with smaller or poorly defined lumina. Semi-automated quantification of histomorpholical features are shown: green outlines are closed pseudoluminal structures and red outlines are open pseudoluminal structures. Schematic illustrations show the characteristic glandular architecture in both groups, highlighting the larger pseudoluminal areas in shedders versus in low/non-shedders. Scale bars: 5 mm (overview), 200 µm (magnification). **B)** Quantification of pseudoluminal area normalized to CK-positive epithelial area in shedders (n = 28) and low/non-shedders (n = 21). Lines indicate mean values, *p*-values were calculated using unpaired two-tailed Mann–Whitney U test. **C)** CK IHC illustrating sites of epithelial barrier loss, resulting in direct exposure of pseudoluminal contents to the surrounding stroma. Scale bars: 100 µm. **D)** Semi-automated quantification of normalized counts of open pseudoluminal structures per shedding group. Lines indicate mean values, *p*-values were calculated using unpaired two-tailed Mann–Whitney U test. **E)** Quantification of pathologist-based scoring of open pseudoluminal structures. Bars show relative distribution per shedding group. *P*-values were calculated from raw patient counts (not shown) using Fisher’s exact test.

At many pseudoluminal sites, cytokeratin (CK) immunostaining revealed partial epithelial barrier loss, resulting in direct contact between pseudoluminal contents and the surrounding stroma (***Fig. 2C***). Such breaches were significantly more frequent in shedders, demonstrated both by semi-automated image analysis (median 0.21 vs. 0.084, *p*=0.0044, ***Fig. 2D***) and visual assessment by trained pathologists (*p*=0.001, ***Fig. 2E***), providing potential conduits for DNA escape into the extracellular space and circulation. Interestingly, both the extent of necrosis, quantified as pseudoluminal-to-epithelial area, and the number of epithelial ruptures showed significant positive correlation with recovery rates of mutations in plasma (***Extended Data Fig. 2***). Detailed spatial IHC profiling of pseudoluminal structures revealed that sites of epithelial barrier loss were enriched for CD68⁺ macrophages (*p*=0.0004), while MPO⁺ neutrophils were significantly increased within pseudoluminal space of shedders (*p*=0.007) (***Fig. 3A-E***). Enlarged, phagocytic CD68⁺ macrophages were also observed within pseudoluminas (***Fig. 3E***).

**Figure 3.**
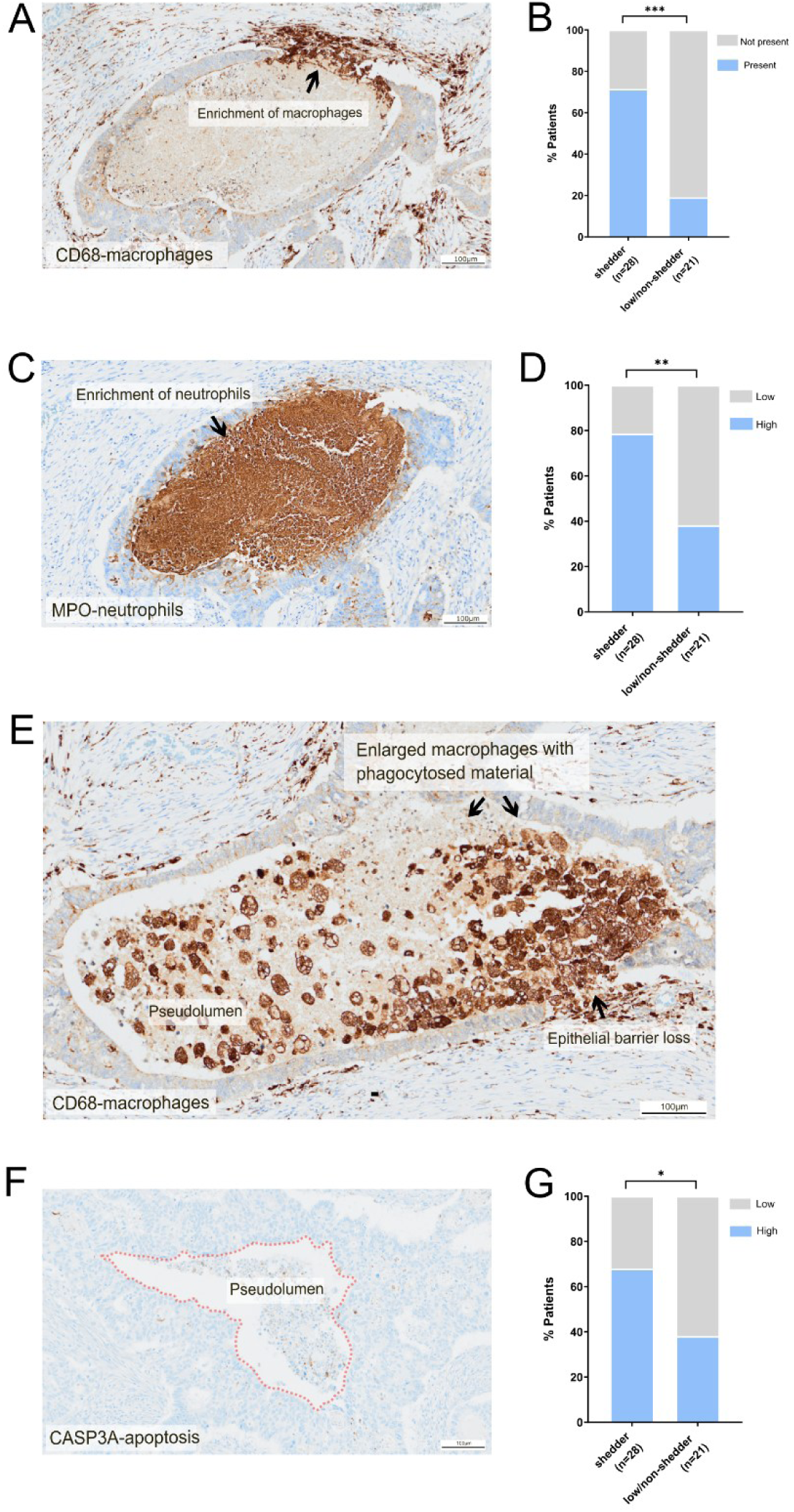
Immune cell enrichment and IHC CASP3A-apotosis quantification at pseudoluminal structures in ctDNA shedder. **A-B**) CD68-macrophage immunohistochemistry (IHC) depicting macrophage influx at sites of epithelial barrier loss in open pseudoluminal structures and corresponding pathologist-based quantification. Bars show relative distribution per shedding group. *P*-values were calculated from raw patient counts (not shown) using Fisher’s exact test. **C-D**) Presence of neutrophils within open pseudoluminal spaces depicted by MPO-neutrophil IHC and correspondent pathologist-based quantification. Bars show relative distribution per shedding group. *P*-values were calculated from raw patient counts (not shown) using Fisher’s exact test. **E**) Infiltration of macrophages at sites of epithelial barrier loss and enlarged macrophages containing phagocytosed cellular debris within pseudoluminal compartments depicted by CD68-macrophage IHC. **F-G**) CASP3A-apoptosis IHC and corresponding pathologist-based quantification within pseudoluminal structures. Bars show relative distribution per shedding group. *P*-values were calculated from raw patient counts (not shown) using Fisher’s exact test.

To further investigate the contribution of apoptosis to ctDNA release, we performed IHC staining for cleaved caspase-3 (CASP3A). While CASP3A positivity across the overall tumor area did not differ between groups (***Supplementary Fig. 3***), analysis restricted to pseudoluminar compartments revealed significantly higher apoptotic activity in ctDNA shedders compared to low/non-shedders (*p*=0.048, ***Fig. 3F-G***). This localized apoptotic enrichment within glandular structures, together with the presence of extensive necrotic pseudolumina, suggests that multiple, spatially distinct modes of cell death contribute to ctDNA shedding.

When analyzing distinct tumor compartments, including the center and invasion front, no significant differences were observed between shedding groups in the densities of MPO⁺ neutrophils, CD68⁺ macrophages, CD4⁺ helper T cells, or CD8⁺ cytotoxic T cells, nor in Ki67 activity (***Supplementary Fig. 3***).

Taken together, ctDNA shedders exhibit a distinct histopathological phenotype characterized by extensive necrotic pseudoluminal architecture, frequent epithelial barrier loss exposing luminal debris to the stroma, localized enrichment of macrophages and neutrophils in and around these sites, and compartment-specific increases in apoptosis. This combination of structural disintegration, immune-mediated remodeling, and spatially confined cell death likely provides multiple coordinated pathways through which tumor-derived DNA can enter the circulation.

### Spatial transcriptomics links necrotic pseudoluminal architecture to myeloid infiltration and stress-associated malignant program

Next, we sought to molecularly characterize the microanatomical regions associated with ctDNA shedding, specifically, pseudoluminal structures and areas of epithelial barrier loss, using Xenium, an image-based spatial transcriptomics platform. To this end, we profiled tumor tissue from two shedders (#218S, #260S) and two low/non-shedders (#234L/NS, #253L/NS) using the Xenium Human Colon Gene Expression Panel (352 genes; ***Fig. 4A***). Following cell segmentation and classification, malignant epithelial and non-malignant cell populations were identified, encompassing a total of 25 distinct cell types across all samples (***Fig. 4B, Extended Data Fig. 3***). As expected, malignant epithelial cells clustered primarily by patient. For example, all neoplastic cells from patient #218S were more similar to each other than to tumor cells from patient #260S, indicating pronounced inter-tumoral heterogeneity in malignant cell programs. In contrast, stromal and immune cell populations were consistently represented across all samples suggesting a relatively conserved composition of the tumor microenvironment despite variability in the neoplastic compartment (***Fig. 4B***).

**Figure 4.**
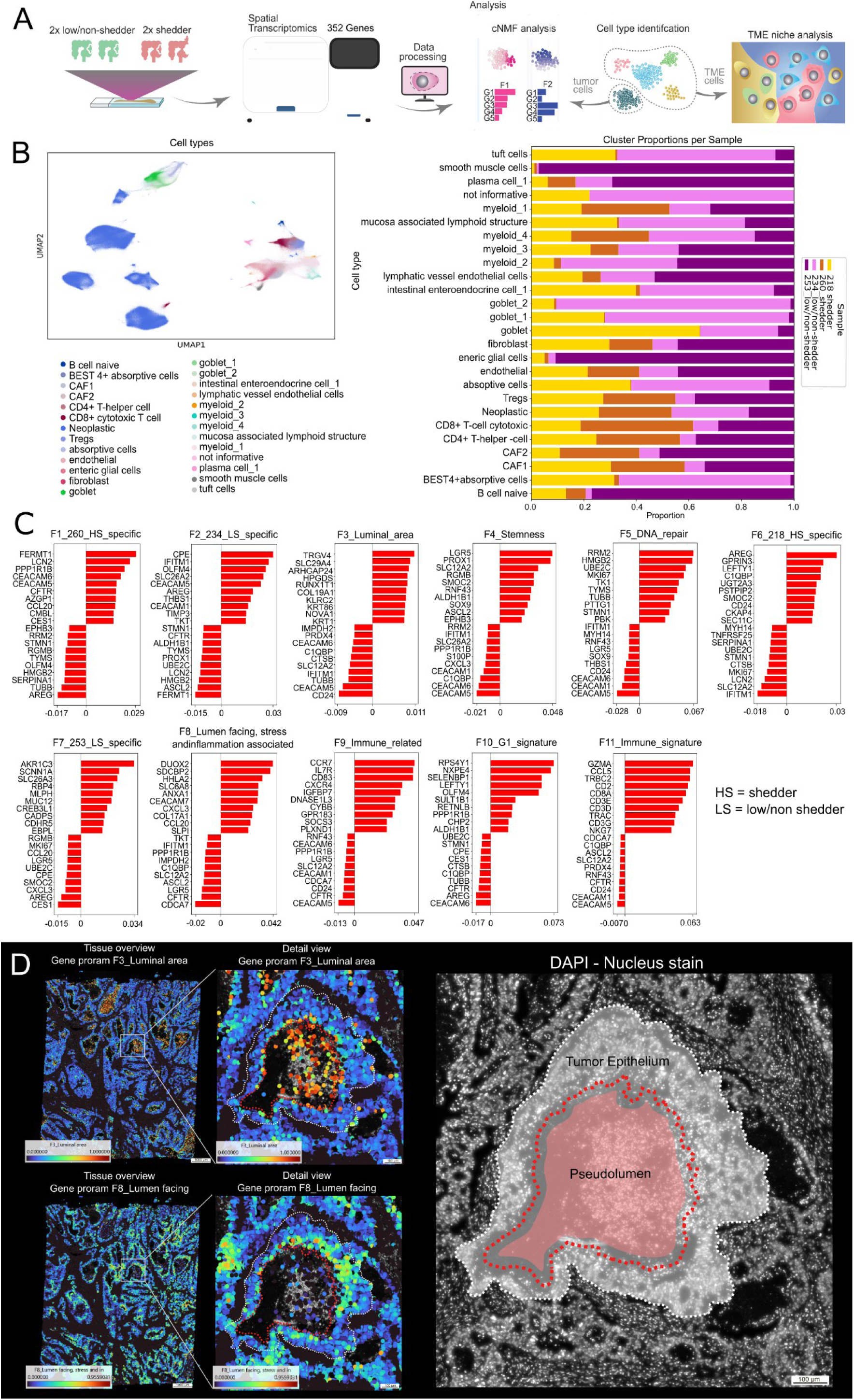
Spatial transcriptomics defines malignant programs and cellular composition in ctDNA shedders versus low/non-shedders. **A)** Overview of the spatial transcriptomics workflow applied to two ctDNA shedders and two low/non-shedders using the Xenium Human Colon Gene Expression Panel (352 genes). Analysis steps included cell type identification, cNMF analysis discovery, and tumor microenvironment (TME) niche analysis. **B)** UMAP of all segmented cells across samples, annotated into 25 cell types. The bar plot shows relative proportions of each cell type per sample. **C)** Eleven malignant epithelial gene programs identified by cNMF, shown with the top contributing genes per program. Programs F3 (“Luminal_area”) and F8 (“Lumen facing, stress and inflammation associated”) were specifically enriched in ctDNA shedders. **D)** Spatial representation of programs F3 and F8. Heatmaps show enrichment of F3 within necrotic pseudoluminal regions and F8 in malignant cells facing the pseudolumen. Right, DAPI reference image with pseudoluminal area and epithelial boundaries highlighted. Data represent individual tissue sections from 4 patients. Scale bars, 100–1000 µm as indicated.

We next investigated whether malignant tumor cells of shedders and low/non-shedder exhibited distinct transcriptional activity states defined as coordinated gene programs reflecting specific cellular functions. Consensus non-negative matrix factorization (cNMF) (26) of malignant transcriptomes identified 11 such gene programs (***Fig. 4C***), of which two programs F3 and F8 were enriched at pseudoluminal structures of shedders. Program F3 (Luminal area) was characterized by high expression of *TRGV4, SLC29A4, ARHGAP24*, *HPGDS, KRT86*, and low expression of *CD24, CEACAM5*, and *TUBB*. These genes are largely associated with immune-related signaling, cell-cell interactions, cytoskeletal remodeling, or cell stress, consistent with dying or degenerating epithelial cell. These cells localized within pseudolumina mirroring the extensive necrosis observed histologically (***Fig. 4D***). Program F8 (Lumen-facing stress/inflammation) marked malignant cells lining the pseudolumen, displaying elevated expression of *DUOX2, SDCBP2, HHLA2, SLC6A8*, and *ANXA1*, and low expression of *CDCA7, CFTR, LGR5, ASCL2, SLC12A2,* and *C1QBP.* These genes align with oxidative stress, cytokine signaling, innate immune activation, or loss of stemness, consistent with cells lining necrotic pseudolumina under inflammatory pressure and are associated with invasion and poor-prognosis-related signatures (***Fig. 4D***) (27).

To further assess the tumor microenvironment (TME) architecture, we applied the BANKSY algorithm, which identified 23 spatially organized cellular niches (***Fig. 5A***). These niches represent tissue areas with a similar cellular and molecular composition, typically linked to coordinated cellular functions. An immune-luminal niche was spatially overlapping with pseudoluminal structures (***Fig. 5B***), and cells attributed to this niche were predominantly found in the two shedder samples (33% in #218S and 58% in #260S versus 6% and 3% in the two low/non-shedders of all cells of the immune-luminal niche). This niche consisted of endothelial cells and myeloid cells located within pseudolumina and at the periphery. Given the high heterogeneity of myeloid populations, we further subclustered these cells, identifying 12 transcriptionally distinct myeloid subtypes (***Fig. 5C, Extended Data Fig. 4***).

**Figure 5.**
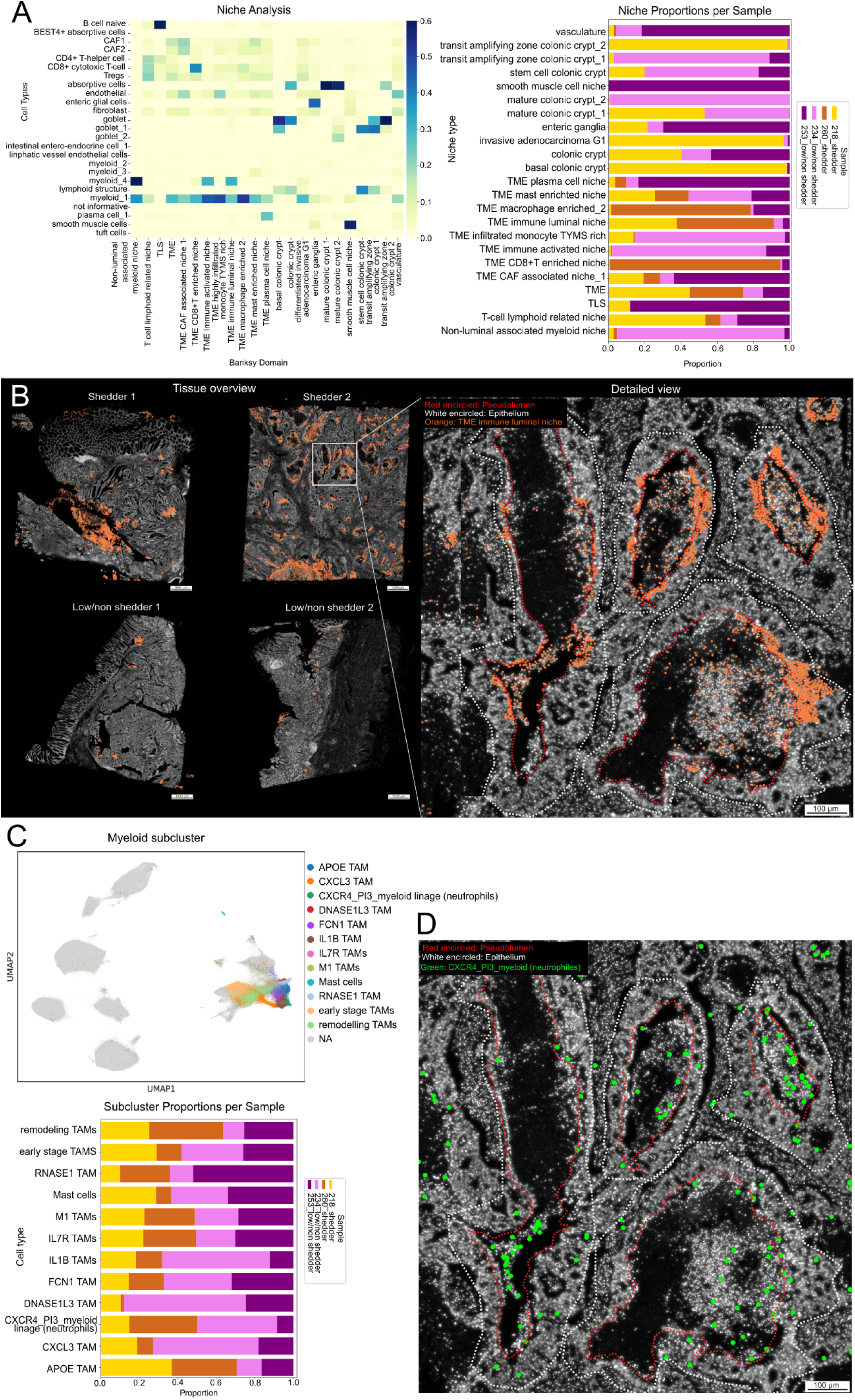
Spatial niche analysis identifies a luminal–interface niche enriched in myeloid populations in ctDNA shedders. **A)** BANKSY niche analysis resolved 23 microenvironmental niches across all samples. The heatmap shows niche-cell type associations. The bar plots illustrate niche distributions per tumor. A distinct luminal-interface niche, characterized by myeloid populations, was higher in ctDNA shedders compared with low/non-shedders. **B)** Spatial mapping of the luminal–interface niche across tissue sections. In shedders, niche cells cluster prominently around pseudoluminal structures, whereas it is only minimally represented in low/non-shedders. The detailed view highlights niche cells located at the epithelial-pseudoluminal interface. **C)** UMAP-based myeloid subclustering identifies 12 transcriptionally distinct states. The bar plot show that several inflammatory and neutrophil-like subclusters are enriched in shedders. **D)** Spatial localization of CXCR4⁺ PI3⁺ myeloid cells (green), demonstrating their accumulation at pseudoluminal regions in shedders. Scale bars: 100–1000 μm as indicated. One Xenium section per tumor (n = 4).

Among these, a *CXCR4+ PI3+* myeloid lineage subtype, consistent with neutrophil-like cells, was specifically enriched within pseudoluminal structures, suggesting a key role in shaping pseudoluminal composition and microenvironmental organization (***Fig. 5D***). Taken together, spatial profiling confirmed that pseudoluminal structures in ctDNA shedders are enriched for myeloid cells, necrosis, and stress-related malignant programs, linking tissue architecture to ctDNA release.

### Spatial mutation detection reveals ctDNA-associated mutations within pseudoluminal structures

To determine whether plasma-detectable tumor mutations are spatially localized within pseudoluminal regions, we applied padlock probe-based *in situ* sequencing (ISS), which allows direct visualization of mutant (MT) and wild-type (WT) alleles in FFPE tissue sections (28,29). We analyzed three shedder CRC samples with tumor-derived mutations detected in matched preoperative plasma (range 10-39 mutations per case) (***Extended Data Table 1***). Strikingly, all plasma-detectable mutations examined were spatially confined in the neoplastic epithelium and the pseudoluminal contents, with 15/15 (100%) detected in sample #260S, 10/10 (100%) in sample #218S, and 39/39 (100%) in sample #223 (***Fig. 6, Extended Data Fig. 5-7***). This spatial co-localization supports a model in which necrosis, apoptosis, epithelial barrier loss, and immune infiltration within pseudoluminal structures create a permissive microenvironment for tumor DNA release into the circulation.

**Figure 6.**
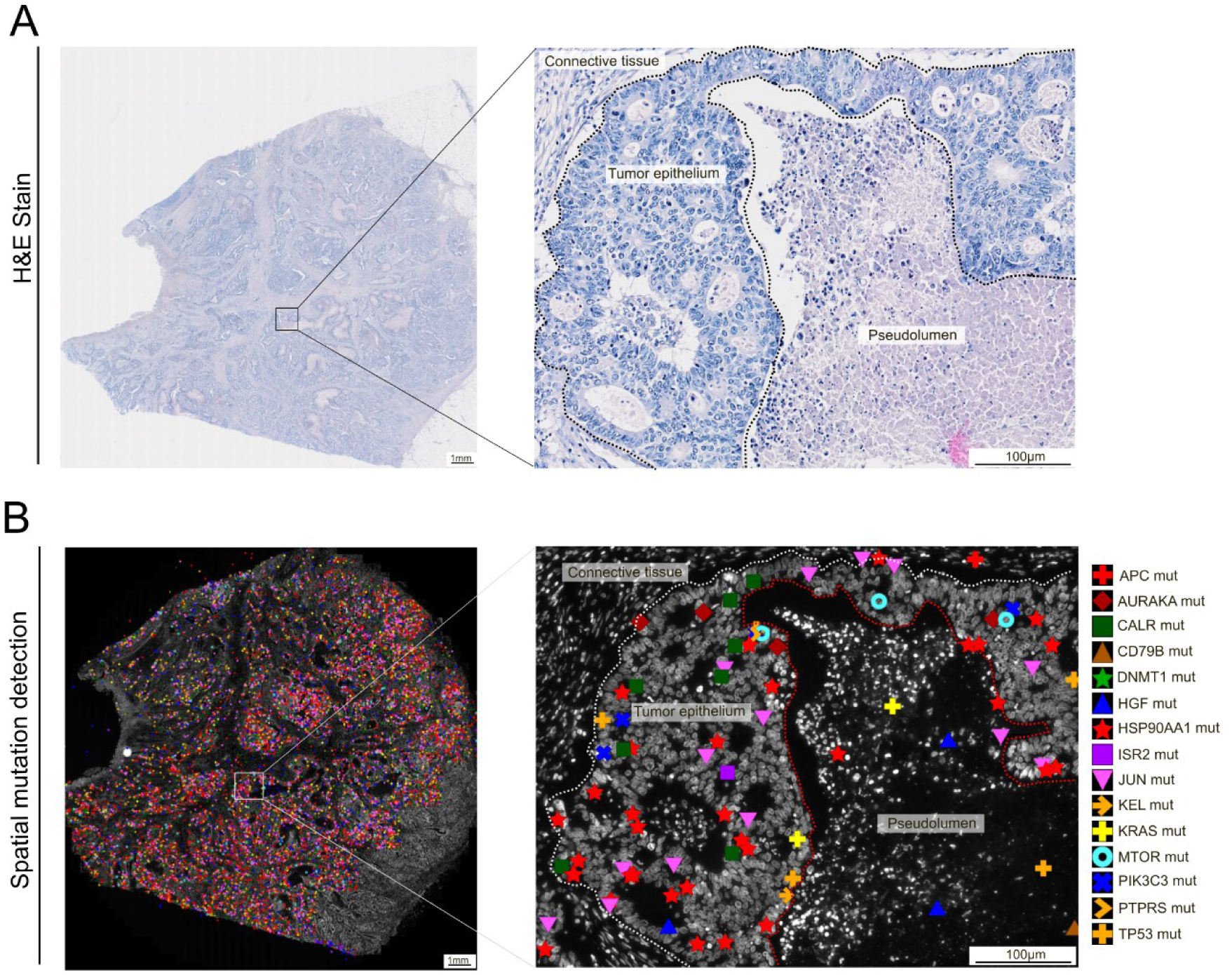
Tumor-derived mutations are spatially detected within pseudoluminal structures and surrounding epithelial tissue. **A)** Histological overview of H&E-stained colorectal tumor tissue from a ctDNA-shedding case (260_shedder). **B)** Spatial *in situ* sequencing (ISS) mapping of somatic mutations, shown as color-coded symbols for individual mutation variants. Mutant allele signals localize within pseudoluminal areas and the surrounding neoplastic epithelium. Scale bars: 5 mm (overview), 100 μm (magnified view).

## Discussion

While tumor size and disease stage have been widely considered as primary determinants of ctDNA, accumulating evidence suggests that biological and microenvironmental factors also play critical roles in governing ctDNA release into circulation (30). In this exploratory study, integrated tumor-plasma sequencing with histopathology and spatial transcriptomics revealed a distinct phenotype in localized CRC ctDNA shedders, marked by necrotic pseudolumina, epithelial barrier loss, and dense myeloid cell infiltration. These features converged into microanatomical niches that concentrate tumor-derived DNA and may represent key sources of circulating ctDNA.

Consistent with previous reports, detectability and recovery rates in our cohort of localized CRC patients increased with advancing disease stage and tumor size, confirming tumor burden as well as the degree of invasion as an important determinant of ctDNA shedding in treatment-naïve CRC (4,31). However, as previously reported pronounced inter-patient variability persist within each stage (32), which indicates that tumor burden alone does not fully explain the magnitude of ctDNA release. Recently, Andersen et al. reported tumor size and proliferative capacity as key contributors to shedding in CRC but emphasized the need for histology-specific metrics (33). Contrary to their observations, we did not identify significant associations between ctDNA recovery and TMB, MSI status, or proliferative activity. Their larger cohort and use of a more analytically sensitive assay may have increased power to detect such relationships. While bespoke assays, as used by Andersen and colleagues, preselect a limited set of clonal mutations and can therefore achieve very high analytical sensitivity, such designs risk biasing conclusions about ctDNA release mechanisms toward those specific variants. In contrast, our approach of using the same panel for plasma and tissue avoids these constraints, enabling unbiased quantification of ctDNA recovery rates. Moreover, Ki-67 reflects current proliferation, whereas ctDNA release is driven primarily by past cell death, microenvironmental stress, and mechanisms of DNA release or clearance, processes that may not correlate directly with proliferative indices. Furthermore, given the inherently high proliferative rate of CRC, subtle Ki-67 differences may be difficult to resolve, making microenvironment-driven shedding more influential.

Our findings instead implicate tumor-tissue architecture and microenvironmental dynamics as primary modulators of ctDNA release, providing a mechanistic explanation for the heterogeneity in shedding in early-stage neoplasms. Our quantitative histomorphological analysis revealed that ctDNA shedders exhibit strikingly expanded pseudoluminal spaces filled with necrotic detritus. When normalized to the epithelial area, pseudoluminal expansion was among the strongest histological correlates of ctDNA recovery. CK immunostaining frequently revealed loss of epithelial barrier integrity exposing necrotic lumina directly to the stroma. The combination of structural breakdown and stromal access provides a plausible pathway for tumor DNA to enter interstitial fluids and ultimately circulation. Both necrosis and epithelial rupture correlated positively with the fraction of tumor variants recovered in plasma, linking tissue architecture to ctDNA release.

Necrosis is a well-established source of cfDNA in cancer (25,34), with associations reported in advanced CRC (35,36), lung (37), and pancreatic cancers (38). In contrast to apoptosis, necrotic cell death generates high local concentrations of larger DNA fragments through uncontrolled genomic release and reduced clearance. However, DNA released through non-apoptotic pathways can be fragmented extracellularly by DNases, producing apoptotic-like size profiles (39). The pseudoluminal architecture observed here provides a distinct morphological niche for such release - confined cavities of dying cells that are directly open to the stroma, likely functioning as local reservoirs from which DNA can diffuse or transit into the bloodstream.

Spatial immunophenotyping revealed that epithelial barrier loss were consistently accompanied by localized innate immune infiltration, dominated by CD68⁺ macrophages and MPO⁺ neutrophils. Enlarged, phagocytic macrophages were frequently observed within pseudoluminal debris, while neutrophils accumulated especially in and around these necrotic spaces. Neutrophils, particularly those engaged in neutrophil extracellular trap (NET) formation, have been implicated in cfDNA release in cancer (40) and recent studies demonstrate that tissue-infiltrating neutrophils can undergo reverse transendothelial migration, a process in which tissue-infiltrating neutrophils re-enter the circulation through the endothelial barrier (41–43). This could, in theory, facilitate the transport of tumor-derived DNA fragments from the tumor microenvironment towards vascular compartments. Although not yet demonstrated in cancer, this mechanism may represent an additional route for ctDNA dissemination.

Macrophages, which phagocytose necrotic debris, may also release tumor-derived nucleic acids during processing (44,45). Although we did not directly quantify NETosis or phagocytic turnover the strong spatial co-localization of these innate immune cells with sites of epithelial barrier loss supports their functional involvement in ctDNA shedding. Both macrophages and neutrophils can release extracellular traps and nucleases that fragment genomic DNA, potentially influencing the composition and fragment size of ctDNA (46). In contrast, adaptive immune markers (CD4⁺ and CD8⁺ T cells) did not differ between shedders and low/non-shedders, suggesting that ctDNA release is shaped primarily by innate rather than adaptive immune activity.

Although necrosis dominated the pseudoluminal architecture, CAS3A staining showed increased apoptotic activity specifically in the epithelial lining of the pseudoluminal structures in shedders. While apoptotic DNA is typically cleared rapidly (47), apoptotic bodies exposed at sites of epithelial barrier loss may evade clearance and contribute to measurable ctDNA. The coexistence of apoptotic rims and necrotic cores suggests that multiple, spatially distinct cell death pathways operate in parallel, likely increasing both the amount and diversity of DNA released into circulation. Therefore, DNA shedding seem to be not a diffuse process but one that occurs at discrete tissue interfaces where structural disintegration meets immune activity. Spatial transcriptomic profiling provided molecular context for the histopathological findings. Stress-responsive epithelial programs enriched in shedders, marked by upregulation of DUOX2, SDCBP2, and ANXA1 and downregulation of differentiation markers, mirrored the histological features of luminal stress and structural breakdown, implicating these epithelial border zones as active contributors to ctDNA release. BANKSY niche analysis further identified a TME immune luminal niche dominated by myeloid cells, with subclustering revealing a CXCR4⁺/PI3⁺ neutrophil-like population specifically enriched in pseudolumina of shedders. This myeloid-rich compartment spatially overlapped with necrotic debris, consistent with the immune - epithelial interactions described above. Importantly, ISS detected plasma-identified mutations within pseudoluminal necrotic material in all tested shedders, providing direct molecular evidence that these structures act as a reservoir of ctDNA. In contrast to malignant epithelial cells that showed strong inter-patient heterogeneity and clustered by individual rather than shedding status, stromal and immune compartments were relatively conserved across samples. This suggests that shedding-related differences arise from local microenvironmental interactions rather than global tumor cell states. This integrated framework may explain why ctDNA levels vary widely among tumors of similar size and stage and highlights the importance of local tissue architecture in shaping liquid biopsy readouts.

Understanding the spatial origins of ctDNA has important implications for interpreting liquid biopsy results. The strong association between pseudoluminal necrosis and ctDNA shedding suggests that tumors with limited pseudoluminal formation may yield lower ctDNA levels despite substantial disease burden, whereas highly necrotic or inflamed tumors may produce disproportionately high ctDNA signals. Incorporating these tissue-level determinants could improve patient stratification for MRD monitoring and early detection.

Several limitations should be considered. First, the sample size of spatial transcriptomic and ISS analyses was modest, reflecting the technical and material constraints of high-resolution assays in FFPE tissue. Validation in larger, independent cohorts is warranted to confirm the generalizability of the pseudoluminal phenotype. Second, our study focused on resected, treatment-naïve tumors; whether therapy-induced changes, such as increased apoptosis or immune infiltration, modulate these structures and alter ctDNA shedding remains to be determined. Third, although ISS provides direct localization of mutant DNA, it does not distinguish between intracellular and extracellular DNA pools, and additional methods will be required to track the kinetics of DNA release and clearance. Finally, while we used a harmonized tumor-informed assay to ensure unbiased variant detection, future work integrating fragmentomics and methylation profiling could further dissect the molecular signatures of ctDNA originating from specific tissue compartments.

In conclusion, this study provides multi-layered evidence that ctDNA shedding in CRC is governed not only by tumor burden but by specific histopathological and microenvironmental features. ctDNA shedders exhibit a characteristic tissue architecture marked by necrotic pseudoluminal structures, epithelial barrier disruption, and myeloid-rich inflammatory niches. These sites combine structural breakdown, immune activity, and spatially organized cell death to create focal reservoirs of tumor DNA that enter the circulation. By linking tissue-level pathology with plasma-derived molecular signals, our findings establish a mechanistic framework for understanding inter-patient variability in ctDNA detectability and lay the groundwork for future approaches to interpret liquid biopsy results in the context of tumor biology.

## Supporting information

Supplementary Figure 1. Comparison of sequencing quality metrics of FFPE and Fresh Frozen (FF) CRC tissue samples using the TSO500 assay. TSO500 pipel

Supplementary Figure 2. Tumor mutational burden (TMB) does not correlate with variant recovery in plasma. Association of TMB with variant recovery in

Supplementary Figure 3. Immune cell distribution and proliferative capacity at tumor center and invasive front of ctDNA shedders vs. low/non-shedders.

Supplementary Figure 4. TSO500 library input and median exon coverage for cfDNA libraries. A) cfDNA input amount used for TSO500 library preparation i

Extended data table 1: In situ sequencing

## Acknowledgments

In-kind reagent support was provided by Illumina, who had no influence over any aspect of the study design, data analysis or reporting. We sincerely thank all patients who participated in the study.

## Consent for Publication

All authors have revised the manuscript and consented to its publication.

## Data Availability

Sequencing data generated in this study for primary tissue and cfDNA have been deposited in the European Genome-phenome Archive (EGA). Access to datasets is granted to qualified researchers upon reasonable request and completion of Data Access Agreement, subject to approval by the local Data Access Committee. All other data supporting the findings of this study are available within the article and its Supplementary Information files. Processed data objects (e.g., count matrices, spatial annotations, niche classifications, etc.) are provided in the Supplementary Data. Custom code used for *in situ* sequencing or Xenium data processing and downstream analyses is available at: https://github.com/Moldia/HybrISS https://github.com/Moldia/CRC_Shedding_Xenium

## Methods

### Patient recruitment and sample collection

In this prospective study, 59 patients with stage I–III CRC diagnosed following routine colonoscopy were enrolled across multiple regional institutions, including LKH II, Hospital St. John of God, and Elisabethinen Graz. Peripheral blood was collected prior to curative tumor resection into PAXgene Blood ccfDNA tubes (PreAnalytiX) and processed within 1–3 days. Plasma isolation was performed using a standardized double centrifugation protocol (1900 × g for 10 minutes, two cycles) to effectively deplete cellular contaminants and minimize genomic DNA carryover. Resulting plasma fractions were aliquoted in 2 mL portions and stored at −80 °C until DNA extraction.

Tumor material was obtained from both formalin-fixed paraffin-embedded (FFPE) blocks collected postoperatively and fresh tumor tissue harvested intraoperatively and immediately snap-frozen to preserve nucleic acid integrity.

FFPE samples were reviewed by certified pathologists, who identified and macrodissected regions with adequate tumor cellularity for downstream molecular analyses. The study was approved by the Ethics Committee of the Medical University of Graz (32-489 ex 19/20), and written informed consent was obtained from all participants. The study complies with the Declaration of Helsinki and good clinical practice.

### DNA extraction

cfDNA was isolated from 8 ml of plasma using the of the QIAamp Circulating Nucleic Acid Kit (QIAGEN) following the manufacturer’s instructions. Genomic DNA from tumor tissue was extracted either from snap-frozen specimens using the QIAamp DNA Mini Kit (QIAGEN) or from FFPE material using the GeneRead DNA FFPE Kit (QIAGEN), applying standardized deparaffinization and DNA repair procedures. DNA yield and purity were quantified using the Qubit 1X dsDNA High Sensitivity Assay Kit (Invitrogen).

### Sequencing workflow and ctDNA detection strategy

To determine the presence and level of ctDNA in plasma, we applied a matched panel tumor-informed ctDNA detection strategy. To this end, tumor tissue DNA and plasma-derived cfDNA were analyzed using the TruSight Oncology 500 solid or ctDNA assays (both v1, Illumina, San Diego, USA). This hybrid-capture–based panel targets 523 cancer genes and enables the detection of single-nucleotide variants (SNVs), insertions/deletions (indels), copy-number variants (CNVs), gene fusions, as well as microsatellite instability (MSI) and tumor mutational burden (TMB).

For tumor DNA, 200ng of DNA was sheared using a Covaris Ultrasonicator S220 (COVARIS) and processed according to the TSO500 solid assay protocol. For cfDNA libraries a median of 95.5 ng (20.4-263ng) was used (***Supplementary Fig. 4***). Final libraries were quality assessed using Qubit fluorometry (Thermo Fisher Sientific) and Bioanalyzer (Agilent). Libraries were pooled based on mass, re-quantified via qPCR (Agilent) using adapter-specific primers, and sequenced in paired-end mode on either a NextSeq 500 (2 × 101 bp, tumor) or NovaSeq 6000 (2 × 151 bp, cfDNA).

Sequencing data were processed using the DRAGEN TSO500 pipeline (v1.1), applying target median coverage thresholds of >150× for tumor tissue and >1300× for cfDNA libraries. The cfDNA coverage threshold was evaluated only for samples without any detectable tumor-matching variants to avoid penalizing biologically true low-shedding individuals (***Supplementary Fig***. ***4***). Manufacturers’ limits of detection (LOD) were followed for tissue variant calling (minimum VAF >5%). For ctDNA, the standard analytical cutoff (VAF >0.1%) was not applied to variants previously identified in matched tumor tissue in order to enhance sensitivity and mitigate stochastic sampling limitations inherent to low-VAF ctDNA detection. VCF outputs were annotated and filtered using Golden Helix®. Putative germline variants were removed by filtering against population allele frequencies using gnomAD v2, excluding variants with population frequencies inconsistent with somatic origin. In addition, variants detected in both plasma and tumor tissue with VAFs in the range of ∼50% were excluded, as these were considered likely germline rather than somatic events and were incompatible with the expected low ctDNA fractions in early-stage CRC. Variant interpretation followed ACMG/AMP guidelines, with variants classified as pathogenic or likely pathogenic considered driver events. Tumor-derived variants were further categorized as clonal or subclonal, with subclonality defined as variants exhibiting a VAF <20% of the highest somatic VAF within the same tumor sample.

### Definition of ctDNA positivity and shedding classification

Given the inherently low tumor fractions in early-stage CRC and the substantial technical and biological variability affecting very low VAFs, we employed a binary recovery-based approach for ctDNA detection. Only somatic variants identified in the corresponding tumor tissue were considered. For each cfDNA sample, a recovery rate was calculated, defined as the proportion of tumor variants also detected in plasma. A plasma sample was deemed ctDNA-positive if at least one tumor-derived variant was recovered. To capture biological variation in tumor shedding while minimizing misclassification due to stochastic effects at low VAFs, patients were stratified into two groups based on their recovery rates, i) ctDNA shedders with recovery rates ≥20%, and ii) non-/low shedders with recovery rates 0–20%. Low- and non-shedders were pooled into one biologically coherent group, as small differences between zero and a few detected variants are likely dominated by technical and stochastic factors rather than true biological differences.

### Immunohistochemistry

For tissue analysis, one representative FFPE tumor block per patient was selected for morphological and IHC evaluation. Hematoxylin and Eosin (H&E) staining was performed to assess key histopathological features, including ulceration, mucinous components, and solid growth patterns. IHC was conducted on 4 μm sections using DAKO antibodies targeting CD68 (macrophages), MPO (neutrophile granulocytes), CD4 (CD4+T-cell), CD8 (CD8+T-cell), Ki-67 (proliferation), cleaved caspase 3a (apoptosis), and cytokeratin (CKAE1/AE3) (epithelium). Staining was performed on the Dako Omnis automated platform.

Cytokeratin (CK) AE1/AE3 staining was evaluated as a surrogate marker of epithelial barrier integrity, categorized as no rupture (0), minor rupture (1), or major rupture (2).

Image-analysis and scoring was performed by two independent pathologists. Scoring criteria were defined individually for each marker based on expected staining pattern and spatial distribution. Immune markers CD68, MPO, CD4 and CD8 were evaluated using a 4-tier semiquantitative scale (0-3), representing absent (0), low (1), moderate (2), or high (3) immune cell abundance. Ki67 proliferation index was determined as the percentage of positively stained tumor nuclei (0-100%). CASP3A expression was assessed separately for pseudoluminal and tissue compartments, whereas the former was semi-quantified (0-3) and the latter calculated as the percentage of positive cells relative to the area with maximal expression.

For each marker, the mean value of both assessments was calculated and used for statistical analysis and graphical representation. Non-integer mean scores (e.g. 1.5) were rounded down to the next lower integer to ensure consistency in categorical grading. For statistical comparisons 4-tier classifications were dichotomized into low (0-1) and high (2-3) expression groups, whereas 2- and 3-tier classifications were reduced to binary categories of absent (0) versus present (1 and/or 2).

Markers were assessed within different tumoral compartments, encompassing the tumor stromal compartment, pseudoluminal structures, the invasion front, and sites of epithelial barrier loss. Ten cases exhibiting predominant mucinous architecture were excluded from pseudoluminal area quantification and downstream analysis due to lack of clearly discernible epithelial or stromal tissue structures.

### QuPath analysis to quantify pseudoluminal structures

Pseudoluminal structures were defined as cytokeratin-negative regions surrounded by cytokeratin-positive epithelial tumor cells (***Fig. 2).*** Necrotic areas were quantified on cytokeratin IHC stained whole-slide images (WSIs) using a semi-automated segmentation pipeline implemented in *QuPath* (version 0.51, 29203879). Pixel-based threshold classification was applied to the DAB channel at full resolution (0.27 µm/px) to generate two binary masks: (1) cytokeratin-positive epithelial tumor regions (above threshold) and (2) pseudoluminal structures, defined as cytokeratin-negative regions (below threshold) surrounded by cytokeratin-positive tumor cells (***Fig. 2***).

Smoothing sigma values (0–3) and DAB thresholds (0.25–0.45) were optimized for each case. Annotations were generated for regions ≥150 µm², with internal holes ≥700 µm² recorded separately. Output annotations were split into independent objects for further processing.

All QuPath segmented histological samples underwent evaluation by experienced histologists to correct for false positive annotations and annotations of epithelial barrier loss. CK-positive annotations were manually refined to remove incorrect detections. Measurements for both cytokeratin-positive and pseudoluminal areas were exported as annotation files and transferred to Excel. For each patient, total area (µm²) of each structure type was calculated, and the ratio of pseudoluminal area to cytokeratin-positive area was determined.

### Spatial transcriptomics (Xenium Platform)

FFPE tissue sections from two ctDNA shedders (#218S and #260S) and two low/non-shedders (#234L/NS and #253L/NS) were profiled using the 10x Genomics Xenium platform with the Human Colon Gene Expression Panel (352 genes), following manufacturer protocols. Raw transcript-level data were converted to AnnData format via a custom Python workflow, integrating expression matrices, gene annotations, and cell metadata. Nuclei-based segmentation provided by Xenium was used, retaining only transcripts within 1 µm of nuclei and with quality scores > 20.

Cell type identification was performed in *Scanpy* (v1.10.3) using standard filtering, normalization, Leiden clustering, and UMAP embedding, followed by manual annotation based on differential expression, spatial context, and reference datasets. Non-neoplastic cells were reclustered to resolve 24 subtypes, and myeloid populations underwent further subclustering. For each patient sample, the proportion of each cell type, niche or myeloid subcluster was determined by quantifying the number of detected cells/niches of that type and normalizing this value to the total number of detected cells within the respective sample.

Spatial niches were identified with the *Banksy* algorithm, focusing on non-malignant cells to reduce variability, and annotated by cell composition and spatial organization. In neoplastic cells, consensus non-negative matrix factorization (cNMF) identified 11 recurrent gene expression programs, whose usage patterns were mapped in spatial and transcriptional space. We define a niche as a microenvironment that supports a small group of cell types (∼10-100 cells) or cellular states, emphasizing local neighborhoods, rather than larger regions.

For example, a tertiary lymphoid structure (TLS) niche may contain naive B cells, Tregs, CD8+ cytotoxic T cells, and CD4+ helper T cells in close spatial proximity.

For details see ***Supplementary methods*.**

### Statistical analyses

All statistical analyses were performed using GraphPad Prism version 10.3.1 (GraphPad Software, San Diego, USA) or R version 4.3.1 unless otherwise specified. Data were tested for normality using Shapiro-Wilk test. For datasets not following a Gaussian distribution, Mann-Whitney U test was applied, while categorical data were analyzed using Fisher’s exact test. Correlations were assessed using Spearman’s rank correlation coefficient. All p values were two-sided and values < 0.05 were considered statistically significant. Specific tests used for each analysis are indicated in the corresponding Figures.

### Spatial mutation detection by *in situ* sequencing (ISS)

To enable spatially resolved detection of point mutations in tumor tissue of shedders (n=3), we designed a panel of padlock probes (PLPs) targeting both the mutant (MT) and corresponding wild-type (WT) sequences (***Extended Data Table 1***). Reverse transcription oligonucleotides, PLP, bridge-probes and detection probes were designed using CLC Workbench (Version7, QIAGEN) as previously described (29,48) (Design parameters specified 15-nucleotide (nt) target-specific hybridization arms with a melting temperature (Tm) between 65 °C and 75 °C. Each PLP consisted of two target-specific arms flanking i) a 20 nt unique identifier (ID) sequence for molecular barcoding, ii) a 20 nt universal anchor sequence shared across subsets of PLPs, and iii) a 20 nt reporter sequence. All oligonucleotides were synthesized by Integrated DNA Technologies (IDTDNA). PLPs were ordered with a 5′ phosphate modification to enable ligation and were resuspended in TE buffer (10 mM Tris, 0.1 mM EDTA, pH 8.0) at a concentration of 100 μM. Barcode-to-gene assignments were generated using an internal Python script designed to minimize barcode crosstalk and maximize decoding accuracy. A complete list of PLPs, target sequences, and barcode assignments is provided in ***Extended Data Table 1***.

For ISS, FFPE tissue sections (5 µm) were baked (60 °C, 1 h), deparaffinized, rehydrated, and permeabilized in citrate buffer (pH 6, 45 min). After dehydration and drying, hybridization chambers were attached, and in situ reverse transcription was performed (45 °C, 3 h) with target-specific primers. PLP were hybridized (37 °C, 30 min), ligated (45 °C, 45 min), and amplified by overnight rolling circle amplification (RCA) at room temperature. Bridge probes were hybridized (37 °C, 1 h) followed by detection probes (37 °C, 30 min). Background fluorescence was quenched, slides mounted with antifade medium, and imaged in six channels. For sequencing, probes were stripped with 100% formamide between imaging cycles (max 5), and final autofluorescence images were acquired for background correction (29).

Imaging was performed on a Slideview VS200 digital slide scanner (Evident) with an external LED light source (X-Cite Xylis/Novem), a 40× UPLXAPO objective (0.95 NA, Olympus), and an ORCA-Fusion sCMOS camera (2304 × 2304 px, 16-bit, Hamamatsu). DAPI, Cy5, and AF750 were detected using a Semrock pentafilter, and Atto425, Atto488, Cy3, and Texas Red with Spectrasplit filters (Kromnigon), each with specified excitation/emission settings. For each sequencing cycle, slides were imaged in six channels (DAPI, FITC, Cy3, Cy5, Cy7, Texas Red) with extended focus imaging to maximize in-focus signal.

Image analysis combined the Cartana pipeline (was acquired by 10x Genomics), MATLAB tools for alignment/tiling/decoding (https://github.com/Moldia/HybrISS) (49), and a CellProfiler/ImageJ workflow for fluorescence intensity measurement, background subtraction and sequencing cycle alignment (29). The Spot Inspector plugin in TissUUmaps was used for visual validation and spatial mapping (50). Images (.vsi) were converted to .tiff, DAPI channels aligned in MATLAB, and tiles generated for analysis. Channel-specific normalization and autofluorescence subtraction were applied in CellProfiler (51). Decoded transcripts were visualized on DAPI images in TissUUmaps.

## Supplemental Methods

**Notebooks accessible at https://github.com/Moldia/CRC_Shedding_Xenium**

### Preprocessing of Xenium data to AnnData format

Spatial raw transcriptomics data of two high-shedders and two non-shedders were generated using the 10x Genomics Xenium platform with Xenium Human Colon Gene Expression Panel. Raw data was converted to AnnData format using a custom Python workflow for downstream analysis. Essentially, transcript-level data were parsed from the Xenium output folders, and each sample was converted into an AnnData object using a custom function that integrates expression matrices, gene annotations, and cell metadata. Default nuclei staining-based segmentation provided by the platform was used to segment cells, following the best practices described in Marco Salas et al. (52). Transcripts were filtered based on spatial proximity to nuclei (within 1 µm) and quality score (QV > 20) to ensure that only high-confidence, nucleus-associated signals were retained. Filtered AnnData objects for each sample were subsequently concatenated into a combined dataset for integrated analysis.

### Spatial transcriptomics data processing and clustering

Using the combined AnnData object, we first aim to identify cell populations using Scanpy (sc) (version 1.10.3). Standard preprocessing steps were applied, including filtering cells with less than 8 genes and less than 20 counts, followed by library size-based normalization and logarithmic transformation. Next, k-nearest neighbor graph was constructed using (<u>sc.pp</u>.neighbors,k=12). Unsupervised clustering was performed using the Leiden algorithm (sc.tl.leiden), and low-dimensional representation was obtained using UMAP (<u>sc.tl</u>.umap). Next, clusters obtained were annotated based on pre-computed differential gene expression analysis (sc.tl.rank_genes_groups, method = ‘wilcoxon’) and spatial distribution, classifying cells in two main categories: neoplastic and non-neoplastic cells.

### Non-neoplastic cell annotation

Next, **non-neoplastic cells** were reprocessed. Essentially, k-nearest neighbor was recomputed, followed by a second round of Leiden clustering to resolve finer-grained cell subtypes within the non-malignant compartments. Non-neoplastic clusters were then manually annotated based on differential gene expression analysis and spatial distribution of clusters (sc.tl.rank_genes_groups, method = ‘wilcoxon’). The 24 cell types identified were annotated based on (1) their expression profile and (2) location in the tissue, using a single cell RNA reference from Chen et al. 2021 as a reference (<u>cellxgene viewer</u>) and known literature.

### Subclustering of myeloid populations

Due to their relevance in CRC shedding biology, we performed subclustering of the main clustering of myeloid, macrophage DNASE1L3^+^, mast cell_1 and monocyte CCL4^+^ IL1B^+^ populations. Essentially, we re-computed the k-nearest neighbor graph using <u>sc.pp</u>.neighbors (k=7). Unsupervised clustering was performed using the Leiden algorithm (<u>sc.tl</u>.leiden). Resulting clusters were visualized in two dimensions using precomputed UMAP (<u>sc.tl</u>.umap). Subpopulations were manually annotated based on differential gene expression analysis (sc.tl.rank_genes_groups, method = ‘wilcoxon’) and spatial distribution.

### Spatial Domain Detection and Niche Analysis

To identify spatial niches in the tumor microenvironment, we applied the Banksy Python (53) package. Malignant cells exhibited high intra-tumoral heterogeneity, leading to the identification of sample-specific niches when included in niche analysis. To minimize this confounding variability and improve cross-sample comparability, we excluded malignant cells from downstream niche analysis. Instead, we focused exclusively on non-malignant cells, which showed more consistent expression profiles across samples. Consequently, the niches identified in our analysis reflect the composition and organization of the tumor microenvironment (TME). Banksy was run using AGF-based spatial clustering (max_m = 1) with 15 spatial neighbors (k_geom = 15) and a scaled Gaussian weight decay. Clustering was performed using a resolution of 0.9 and 20 principal components. Resulting niches were annotated based on their cell type composition and spatial distribution. Finally, the cell type composition of each niche was visualized as a heatmap using Seaborn, illustrating niche - specific cell type composition.

### cNMF on neoplastic cells to look for gene programs along the dataset

Consensus non-negative matrix factorization (cNMF) was performed on neoplastic cells to identify recurrent transcriptional programs. For this, we employed the workflow from the dylkot/cNMF Python package (26). The model was run with 200 replicates (n_iter=200) over a range of components K = 5 to 15 (components=np.arange(5,16)). A stability and density-based evaluation guided component selection; K = 11 was selected as the optimal number of programs based on k_selection_plot() outputs. The final consensus step was run using a density threshold of 0.10, which defines the minimum similarity for grouping components across replicates (cnmf_obj.consensus(k=11, density_threshold=0.10)). The resulting gene expression programs (GEPs), usage matrix (program contributions per cell), and top genes were stored for further analysis. The usage matrix was merged with the UMAP-embedded AnnData object, and program usage scores were overlaid on the UMAP plot using sc.pl.umap, revealing the spatial and transcriptional structure of neoplastic cells.

**Extended Data Figure 1.**
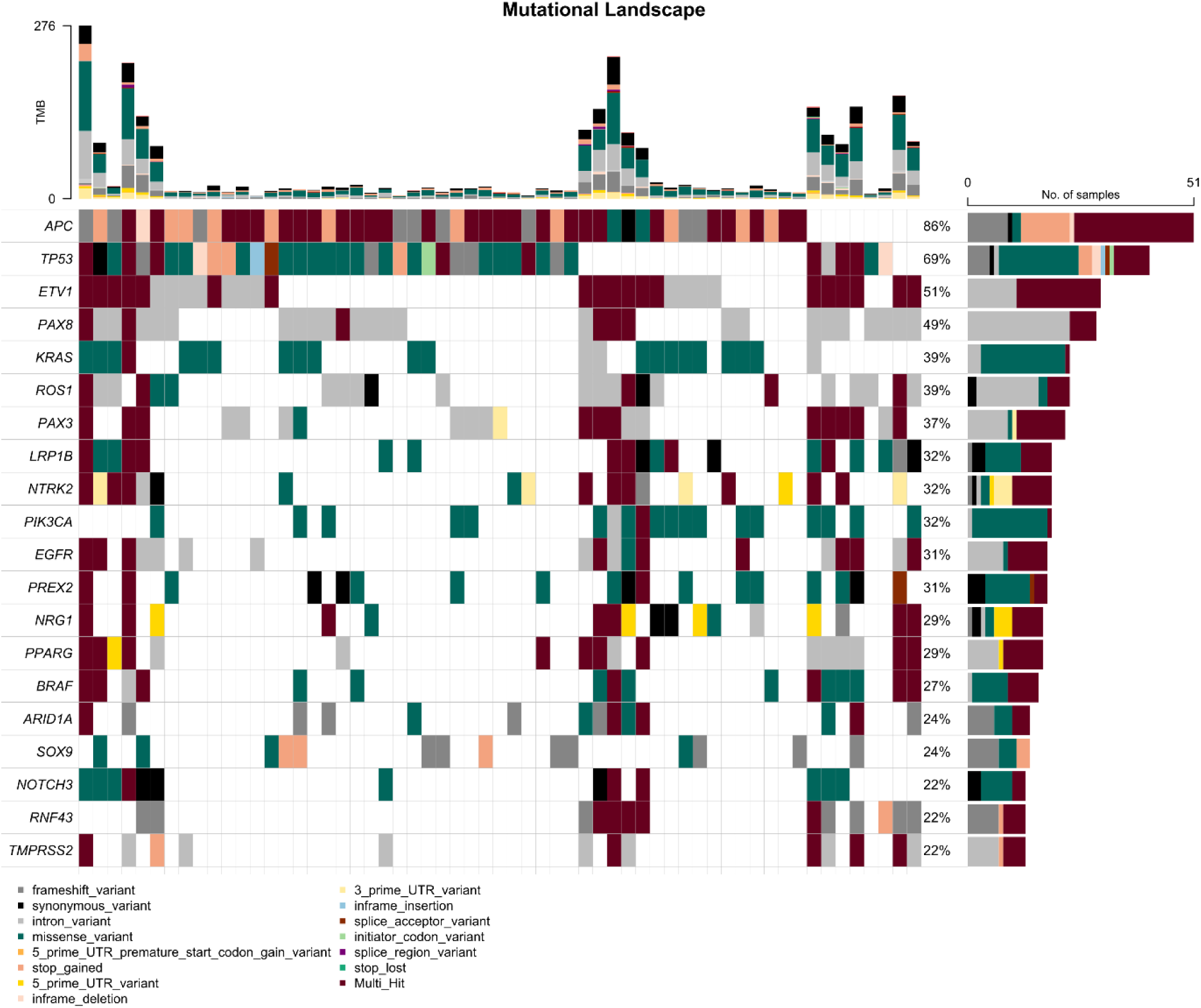
Mutational landscape of primary colorectal cancer tissue. Oncoplot showing the 20 most frequently mutated genes across the cohort (n=59). Columns represent individual patients, rows recurrently mutated genes. Mutation types are color-coded as indicated. Tumor mutational burden (TMB) and percentage of individuals harboring mutations within a specific gene are shown on top or on the right respectively.

**Extended Data Figure 2.**
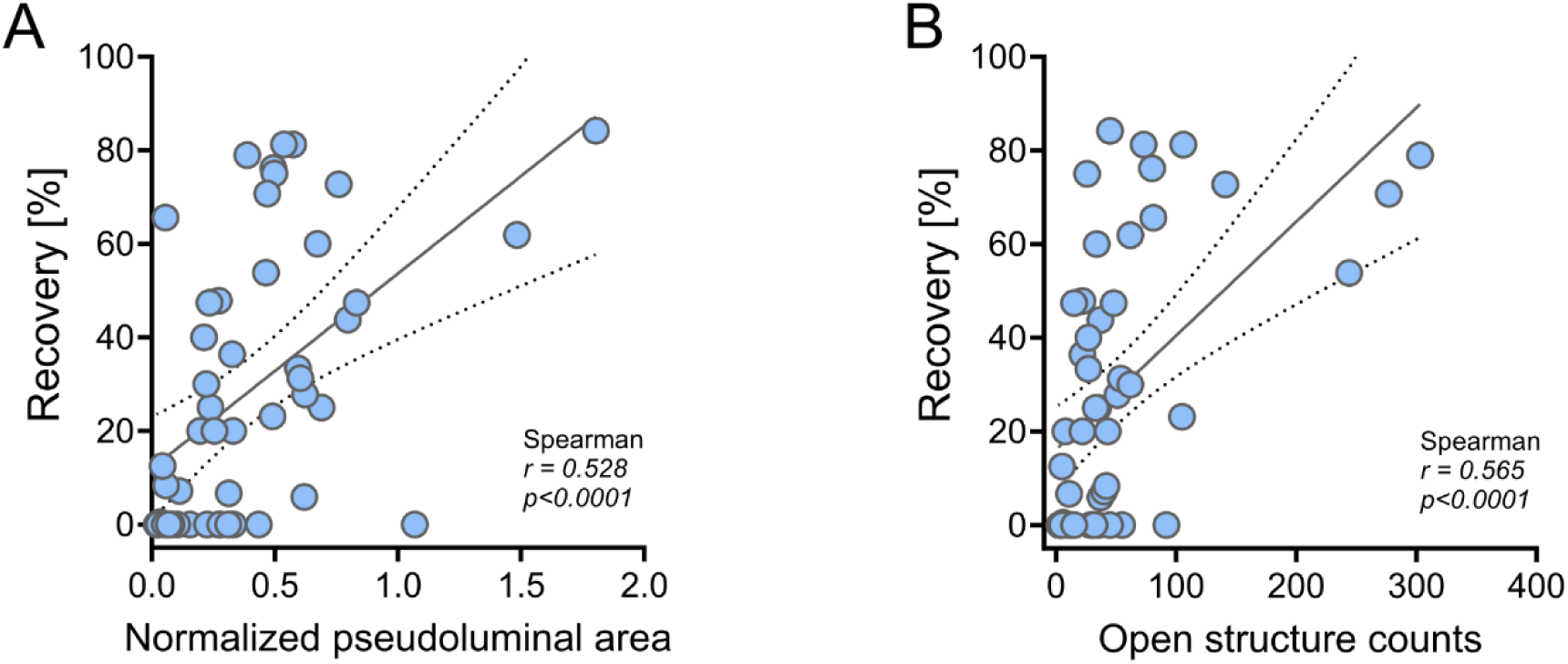
Correlation of tumor tissue variant recovery in plasma with pseudoluminal area and epithelial barrier loss. **A)** Dot-plot showing association of tumor tissue variant recovery in plasma with pseudoluminal area and **B)** epithelial barrier loss depicted by the number of open structure counts. *r*- and *p*-values were calculated using Spearman rank-correlation, a linear regression line with 95% confidence interval is shown for visual reference.

**Extended Data Figure 3.**
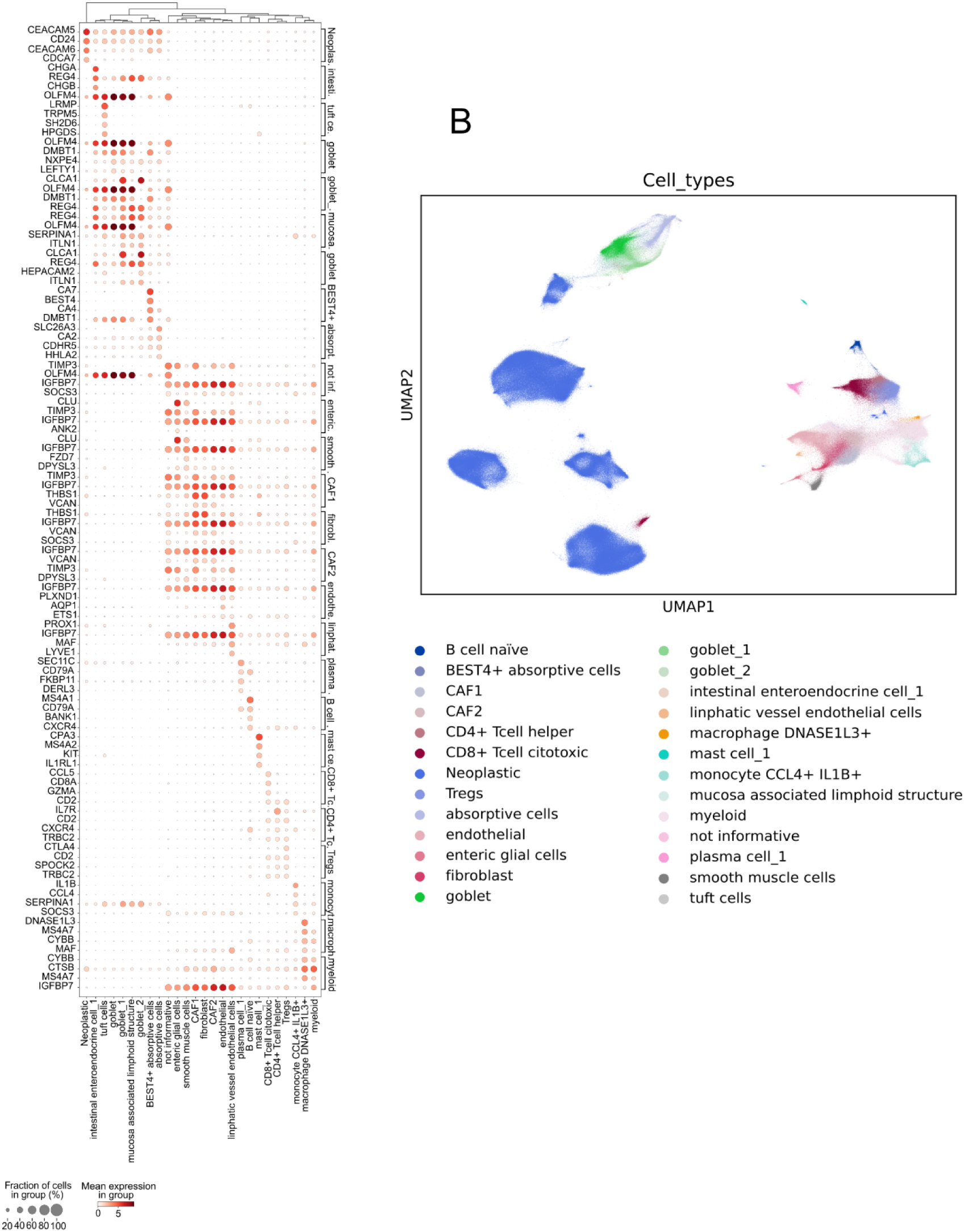
Dot plot of differentially expressed genes across transcriptionally defined cell clusters. **A)** Dot plot showing differentially expressed genes (DEGs) identified by Wilcoxon testing across transcriptionally distinct cell clusters derived from Xenium spatial transcriptomics data. Dot size represents the fraction of cells within each cluster expressing the gene; dot color indicates mean expression level. **B)** UMAP of all identified cell clusters, annotated by cell type.

**Extended Data Figure 4.**
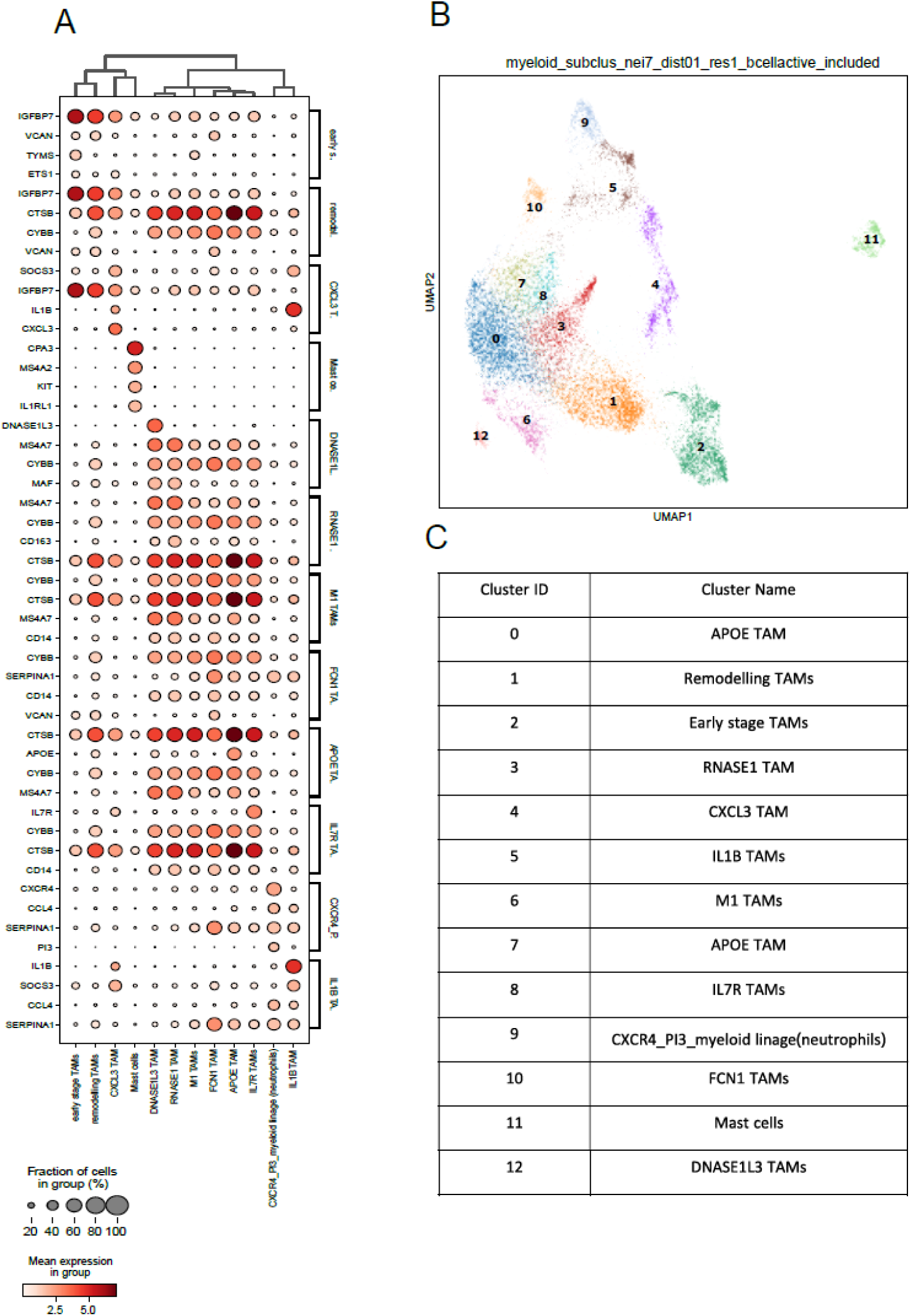
Differentially expressed genes and transcriptional states of myeloid subclusters identified by Xenium data. **A)** Dot plot displaying cluster-specific marker genes across transcriptionally distinct myeloid subclusters derived from Xenium spatial transcriptomics. Dot size represents the fraction of cells expressing each gene within a subcluster, and color intensity reflects mean expression levels. **B)** UMAP of myeloid cells, illustrating 13 transcriptionally defined subclusters. Each subcluster forms a distinct transcriptional neighborhood. **C)** Table listing subcluster identities.

**Extended Data Figure 5.**
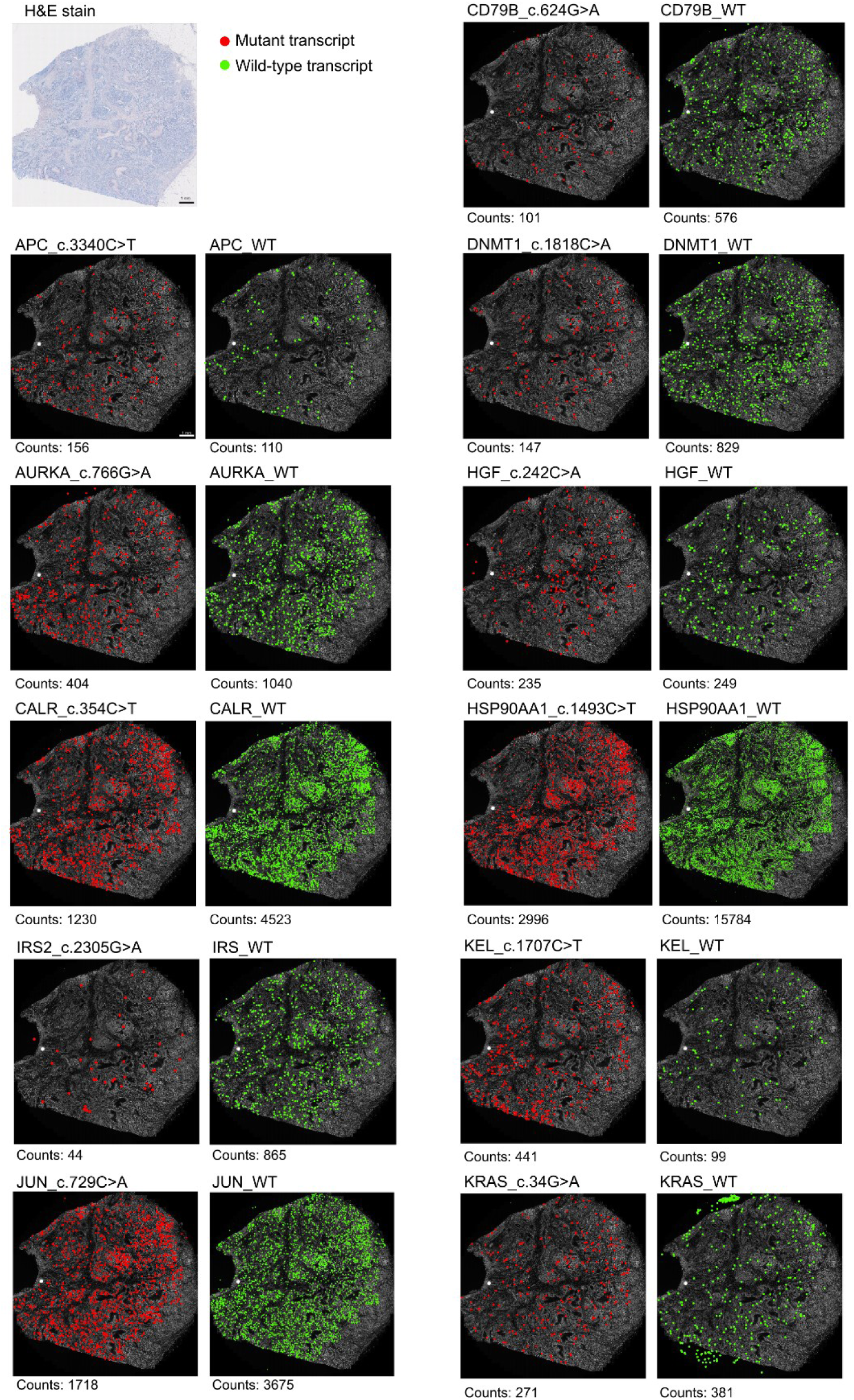
Spatial mutation detection in sample 260_shedder. Spatial *in situ* sequencing (ISS) mapping of tumor-derived mutations in ctDNA shedder sample (260_shedder). Each dot represents the spatial location of an individual detected transcript. Red indicates mutant transcripts and green indicates corresponding wild-type transcripts.

**Extended Data Figure 6.**
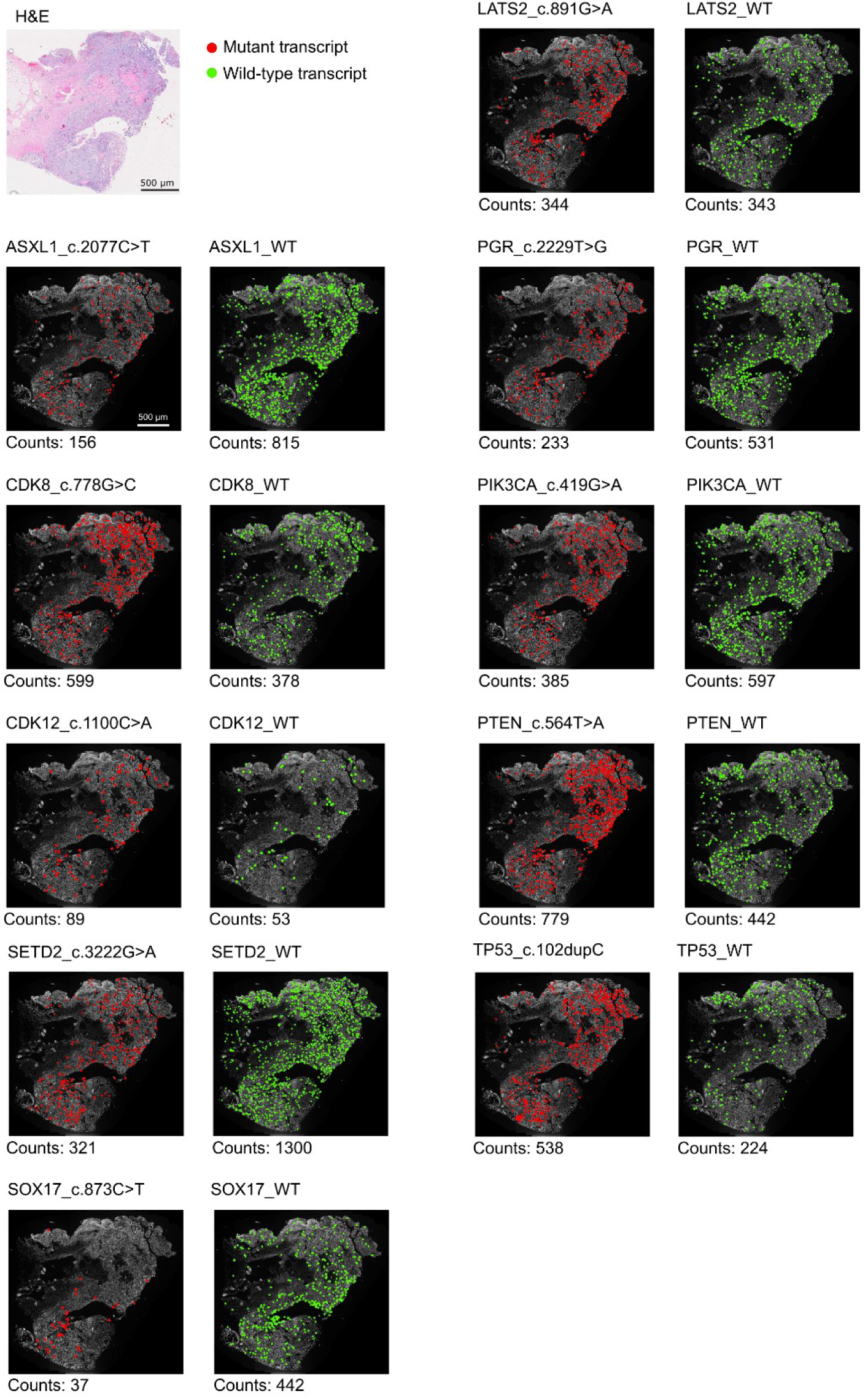
Spatial mutation detection in sample 218_shedder. Spatial in situ sequencing (ISS) mapping of tumor-derived mutations in ctDNA shedder sample (218_shedder). Each dot represents the spatial location of an individual detected transcript. Red indicates mutant transcripts and green indicates corresponding wild-type transcripts.

**Extended Data Figure 7.**
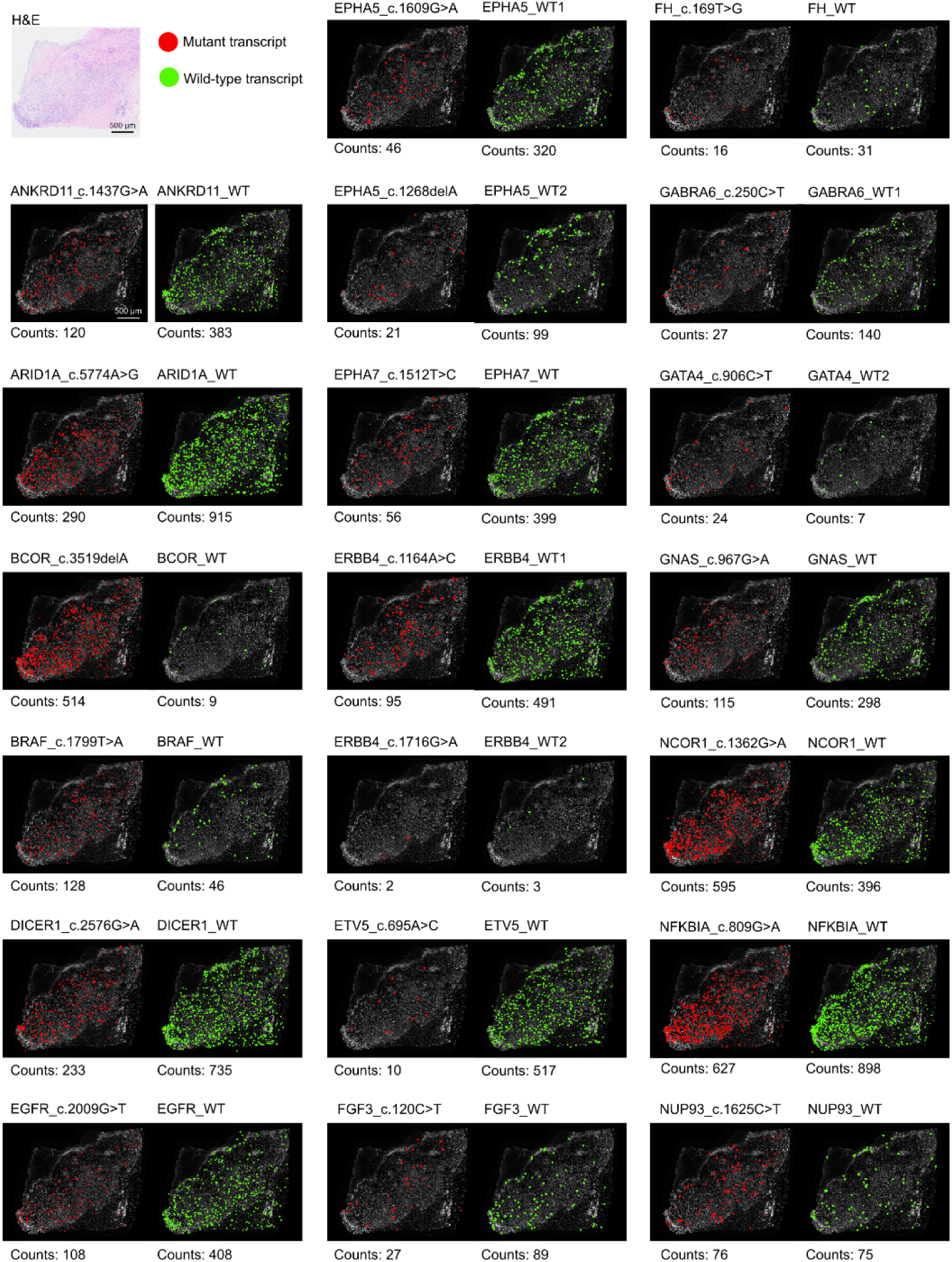
Spatial mutation detection in shedder sample 223_shedder. Spatial in situ sequencing (ISS) mapping of tumor-derived mutations in ctDNA shedder sample (223_shedder). Each dot represents the spatial location of an individual detected transcript. Red indicates mutant transcripts and green indicates corresponding wild-type transcripts.

