## Supplementary figures and images for "Tissue architecture and immune niches govern ctDNA release in colorectal cancer"

### Supplementary Figure 1. Comparison of sequencing quality metrics of FFPE and Fresh Frozen (FF) CRC tissue samples using the TSO500 assay. TSO500 pipel

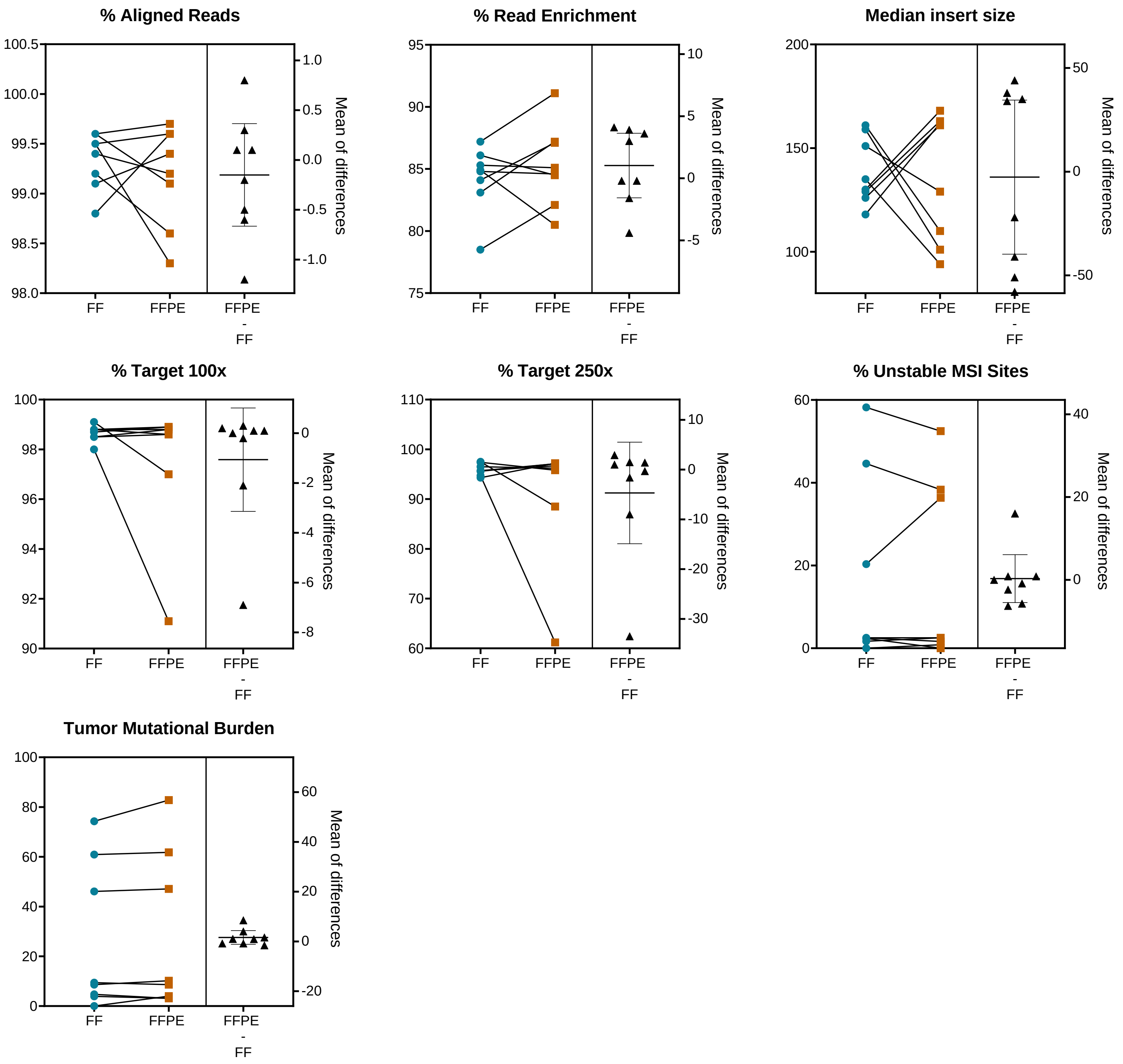

### Supplementary Figure 2. Tumor mutational burden (TMB) does not correlate with variant recovery in plasma. Association of TMB with variant recovery in

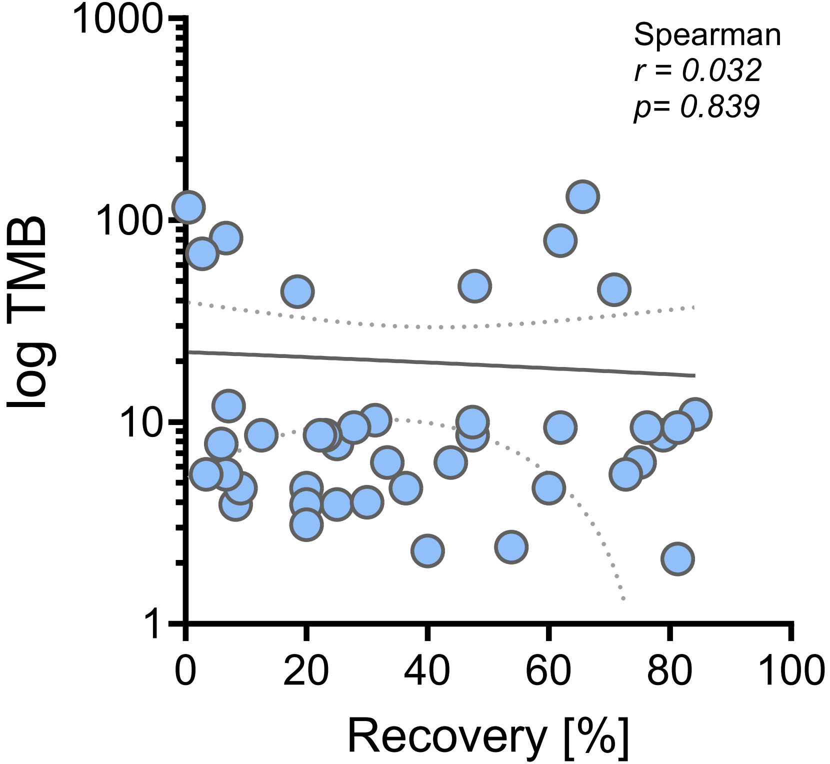

### Supplementary Figure 3. Immune cell distribution and proliferative capacity at tumor center and invasive front of ctDNA shedders vs. low/non-shedders.

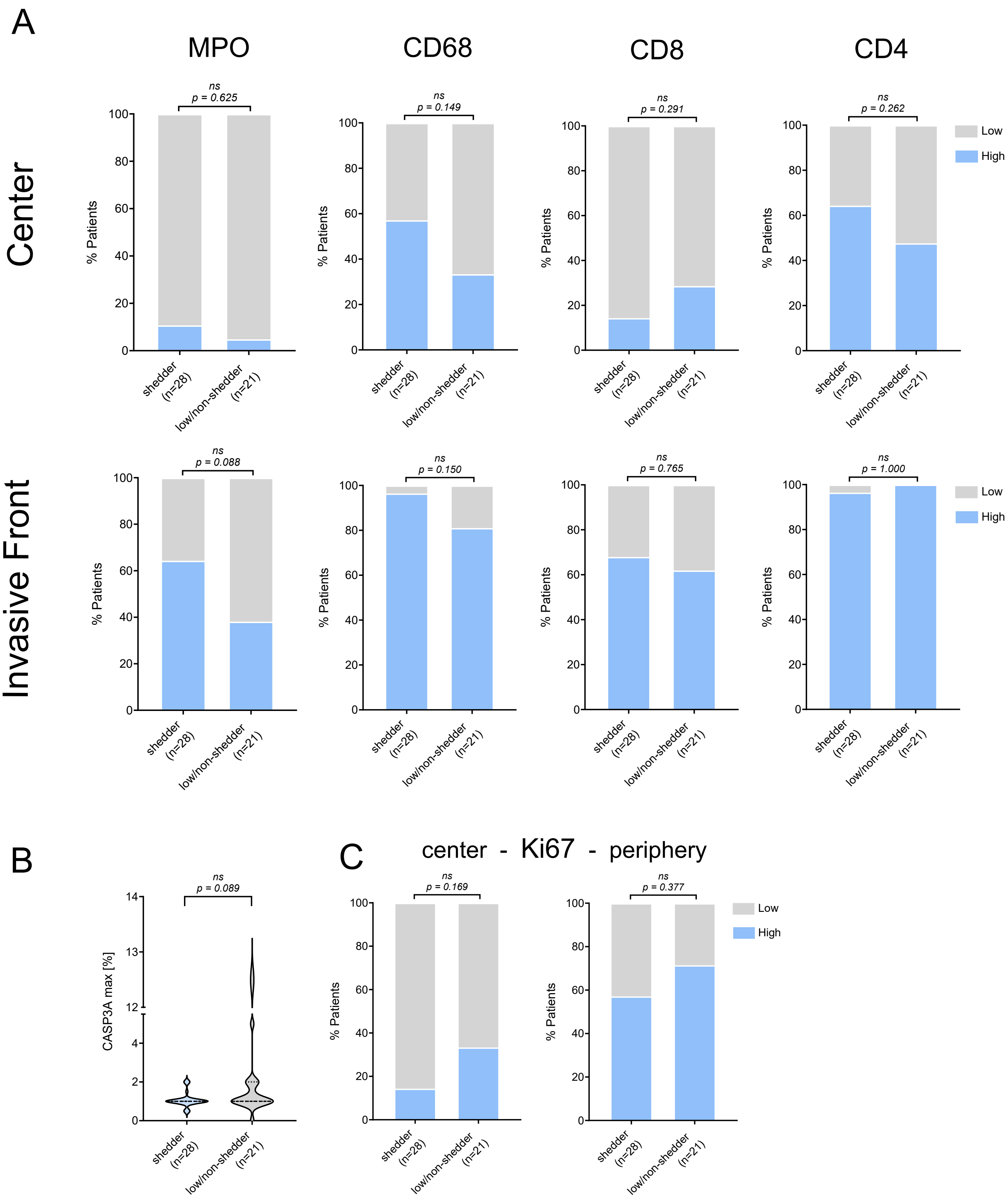

### Supplementary Figure 4. TSO500 library input and median exon coverage for cfDNA libraries. A) cfDNA input amount used for TSO500 library preparation i

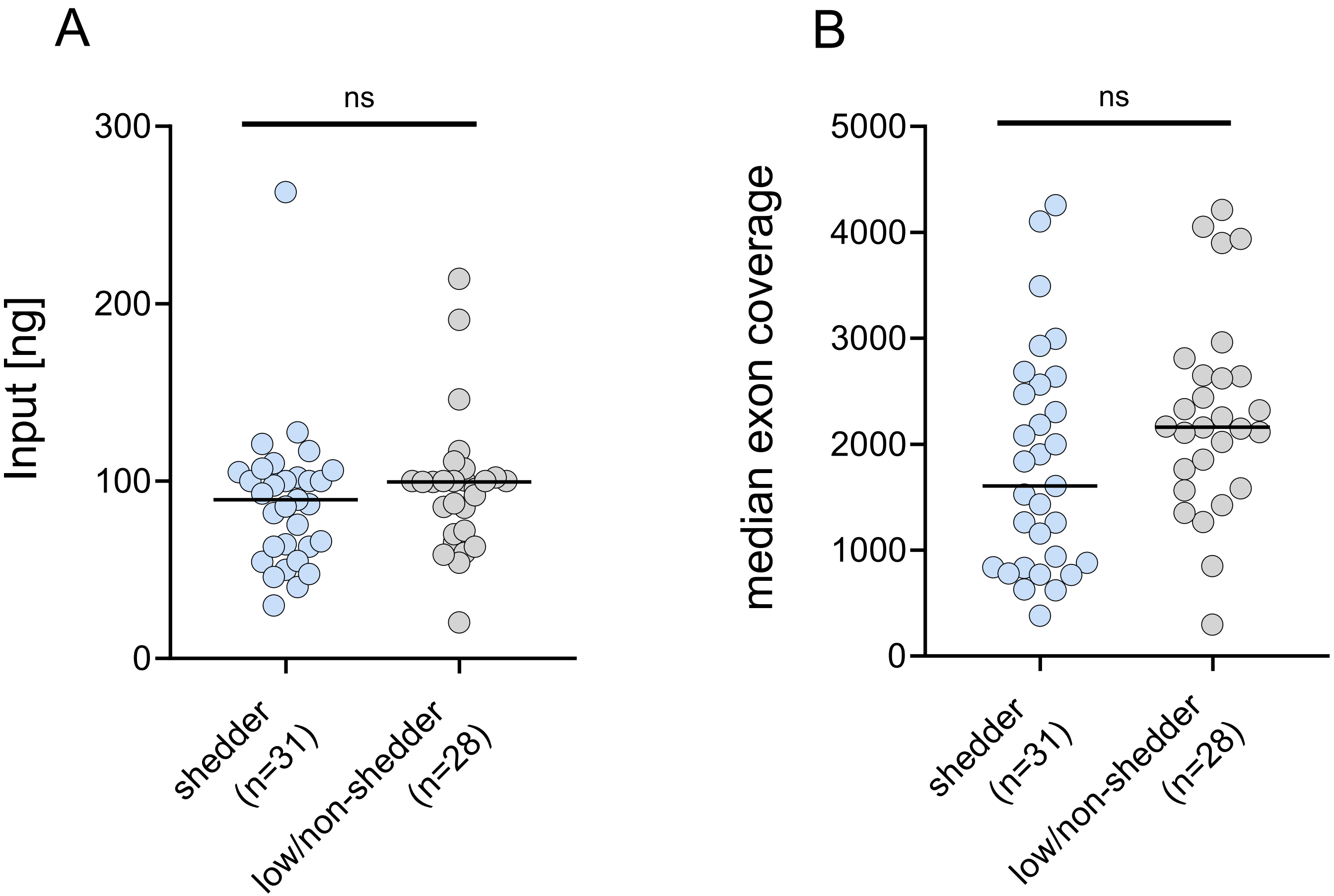
